# Valence ambiguity dynamically shapes striatal dopamine heterogeneity

**DOI:** 10.1101/2024.05.17.594692

**Authors:** Kaisa N. Bornhoft, Julianna Prohofsky, Timothy J. O’Neal, Amy R. Wolff, Benjamin T. Saunders

**Affiliations:** Department of Neuroscience, University of Minnesota; Medical Discovery Team on Addiction, University of Minnesota

## Abstract

Adaptive decision making relies on dynamic updating of learned associations where environmental cues come to predict positive and negatively valenced stimuli, such as food or threat. Flexible cue-guided behaviors depend on a network of brain systems, including dopamine signaling in the striatum, which is critical for learning and maintenance of conditioned behaviors. Critically, it remains unclear how dopamine signaling encodes multi-valent, dynamic learning contexts, where positive and negative associations must be rapidly disambiguated. To understand this, we employed a Pavlovian discrimination paradigm, where cues predicting positive and negative outcomes were intermingled during conditioning sessions, and their meaning was serially reversed across training. We found that rats readily distinguished these cues, and updated their behavior rapidly upon valence reversal. Using fiber photometry, we recorded dopamine signaling in three major striatal subregions −,the dorsolateral striatum (DLS), the nucleus accumbens core, and the nucleus accumbens medial shell - and found heterogeneous responses to positive and negative conditioned cues and their predicted outcomes. Valence ambiguity introduced by cue reversal reshaped striatal dopamine on different timelines: nucleus accumbens core and shell signals updated more readily than those in the DLS. Together, these results suggest that striatal dopamine flexibly encodes multi-valent learning contexts, and these signals are dynamically modulated by changing contingencies to resolve ambiguity about the meaning of environmental cues.

## INTRODUCTION

Adaptive decision making requires rapidly adjusting learned behaviors in response to changing contingencies. Environmental cues guide this process via Pavlovian associations, signaling the valence and identity of future rewards and threats. Understanding how the brain disambiguates complex and dynamic sensory elements to promote flexible behavior is important for insight into a number of psychiatric disorders, including addiction, obsessive compulsive disorder, PTSD, and schizophrenia (Izquierdo and Jentsch, 2012; Leeson et al., 2009; Remijnse et al., 2006; Swainson et al., 2000).

The striatum is a key center for cue-based learning, decision making, and action selection, and disruptions in striatal signaling contribute to behavioral inflexibility (Brown et al., 2011; Castañé et al., 2010; Cox and Witten, 2019; Haluk and Floresco, 2009; Klanker et al., 2013; Ragozzino, 2007). Critically, the striatum is anatomically and functionally heterogeneous (Bakhurin et al., 2016; Barnes et al., 2005; Cardinal et al., 2002; Di Ciano et al., 2001; Everitt and Robbins, 2005; Nestler and Lüscher, 2019; Saunders and Robinson, 2012; Swainson et al., 2000; Yin et al., 2009), and a central component of this is dopamine signaling, where different striatal niches subserve specialized learning-related functions (Brown et al., 2011; Collins and Saunders, 2020; Cox and Witten, 2019; Howe and Dombeck, 2016; Klanker et al., 2017; Mohebi et al., 2024; Parker et al., 2016; Radke et al., 2019; Saunders et al., 2018). While striatal dopamine is classically associated with reward learning and reinforcement (Schultz et al., 1997; Wise, 2004), there is now extensive evidence for a broader role, including encoding negative valence and threat/avoidance conditioning (Badrinarayan et al., 2012; De Jong et al., 2019; Fadok et al., 2009; Lammel et al., 2014; Mccutcheon et al., 2012; Oleson et al., 2012; Pezze and Feldon, 2004; van Elzelingen et al., 2022; Wendler et al., 2014).

Dopamine responses vary widely across striatal regions, and it remains unclear how these signals encode conditioned valence and adapt to situations of valence ambiguity. To explore this, we employed a Pavlovian valence discrimination paradigm (Burgos-Robles et al., 2017), where cues predicting positive and negative outcomes were intermingled during conditioning sessions, and their meaning was serially reversed across training. Rats came to distinguish these cues and updated their behavior rapidly upon valence reversal. Using fiber photometry to measure dopamine signaling in three major striatal subregions - the dorsolateral striatum (DLS), the nucleus accumbens core, and the nucleus accumbens medial shell - we found a triple dissociation in the qualitative pattern of responses to reward and threat conditioned cues and their predicted outcomes. Furthermore, valence ambiguity introduced by cue reversal reshaped striatal dopamine signals on different timelines, with nucleus accumbens core and shell dopamine signals updating faster than those in the DLS. Together, our results indicate that striatal dopamine niches encode different features of positive and negative Pavlovian associations, and these signals are dynamically modulated by changing contingencies to rapidly resolve ambiguity about the meaning of environmental cues.

## RESULTS

### Rats discriminate between intermingled positively and negatively valenced Pavlovian cues

To investigate the effect of intermingled positive and negatively valenced stimuli on conditioned behavior and dopamine signaling in striatal subregions, we used a Pavlovian discrimination task adapted from previous studies (Burgos-Robles et al., 2017). Rats first underwent habituation to three novel auditory cues (high tone, low tone, white noise), followed by an initial learning phase where one conditioned stimulus (CS) was paired with a liquid reward (CS+R), one was paired with a footshock (CS+S) and a third was paired with no outcome (CS-) (**Figure 1A**). To explore the relationship between dopamine signals and dynamic cue valence, the initial learning phase was followed by a reversal phase, where the tones predicting reward and shock delivery were reversed. After further conditioning under the reversed cue conditions, we employed a second reversal phase, where the CS and US pairings were returned to their original contingencies for several sessions (**Figure 1B**). To track cue discrimination, we measured two primary conditioned responses (CR): reward port entries (PEs) and freezing. First we examined the percentage of time spent in the port for the entire 20-s cue period. Rats quickly learned to discriminate between CS types, and this discrimination grew across sessions (**Figure 1C**; session x CS interaction F(8,176)=22.23, p<0.0001).

**Figure 1.**
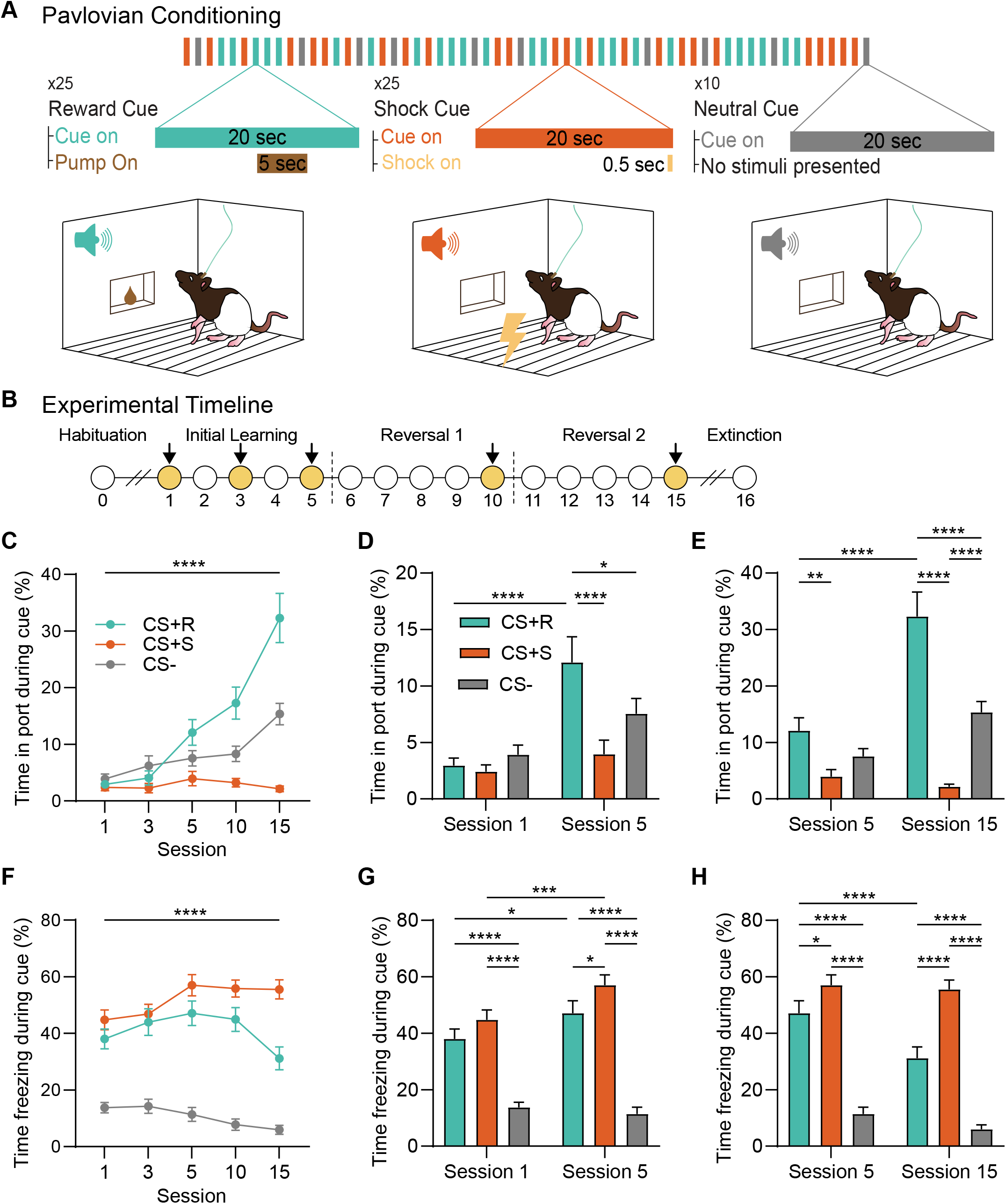
Multivalent Pavlovian cue discrimination task. A Schematic showing a representative session of the task. Shock (CS+S), reward (CS+R), and neutral cue (CS-) trials were intermingled during Pavlovian training. Reward port entries and freezing bouts were the primary measures. Rats (n=23; 11F, 12M) learned to discriminate the cues with appropriate behavioral responses. B Experimental timeline. Primary training data from sessions 1, 3, 5, 10, and 15 are shown in this figure. C Percent time spent in the port during the cues diverged with training, with the most time spent in the port during the CS+R. D Percent time in port during the CS+R increased during initial training, and by session 5 rats discriminated the CS+R from the other cues. E Percent time spent in port during the cues continued to diverge after initial training. F Percent time freezing during the cues diverges across training, with the most time freezing spent during the CS+S. G Percent time freezing during initial training. H Percent time freezing remained divergent after initial training. All bars represent mean ± SEM. ****p<0.0001, ***p<0.001, **p<0.01, *p<0.05.

Time in the port during the cue increased from day 1 to 5 only for the CS+R (**Figure 1D**; multiple comparisons test p<0.0001), and by day 5, time in the port was higher for the CS+R than the CS+S (p<0.0001) and CS(p=0.0240). After the initial training phase (Days 1-5), cue discrimination continued to improve following reversal (**Figure 1E**), as shown by increased time in port from day 5 to 15 for the CS+R (p<0.0001). Time in port for the CS+R on day 15 was higher than for either the CS-or CS+S (both p<0.0001). Notably, time in port also increased from day 5 to 15 for the CS-(p=0.0100), and was higher for the CS-than CS+S on day 15 (p<0.0001), potentially reflecting the learned safety of the CS-period.

Next, we examined the percentage of time rats spent freezing during each cue. Rats quickly learned to discriminate between CS types across sessions (**Figure 1F**; session x CS interaction, F(8,186)=7.071, p<0.0001). By day 5, freezing was highest for the CS+S (**Figure 1G**), relative to the CS+R and CS-(multiple comparisons test, p=0.0109 and p<0.0001, respectively). Across the remaining training phases, cue discrimination improved (**Figure 1H**). Freezing during the CS+S was greater than the other cues (both p<0.0001), and the time spent freezing during the CS+R decreased from day 5 to 15 (p<0.0001). We saw some evidence of CR generalization (**Figure 1F-1H**), as freezing during the CS+R remained higher than the CS-(p<0.0001) across training. We also examined Pavlovian conditioned behaviors with data across sexes (**Suppl. Figure 1**). Overall we found no sex differences in Pavlovian cue discrimination (2-way mixed model ANOVAs, no sex containing interactions and all main effects of sex p>.1) with females (n=11) and males (n=12) exhibiting comparable levels of cue discrimination and conditioned behavior across training.

### Subregion-specific striatal dopamine responses to positively and negatively valenced outcomes

To explore dopamine encoding of cues and unconditioned stimuli in a multi-valent Pavlovian learning context, we recorded dopamine activity in the DLS, NAc core, and NAc shell using fiber photometry with the dopamine sensor dLight 1.3b (**Figure 2A-2B; Suppl Figure 2**). First, we examined dopamine signals in response to each unconditioned stimulus (footshock, reward), finding heterogeneous activity. The dopamine signal in response to reward consumption varied qualitatively across regions: there was a small, slow positive response in the DLS (**Figure 2C**), a larger, fast positive response in the core (**Figure 2D**), and a small, positive response followed by a dip below baseline in the Session shell (Figure 2E). Given this variability, we measured the peak or trough of the reward response, which differed between regions (**Figure 2F**; one-way ANOVA F(2,18)=9.941, p=0.0012). Next, we looked at dopamine responses to shock delivery, which also varied across regions: the DLS had a biphasic response, with a trough during shock followed by a positive peak at shock offset (**Figure 2G**); the core had a positive response during shock (**Figure 2H**); and the shell responded more slowly, with little change during the shock itself but a slow steady increase in dopamine that emerged after shock offset with a mean time to peak of 1.51 s (**Figure 2I**). Correspondingly, the magnitude of the shock response differed across regions (**Figure 2J**; F(2,20)=24.54, p<0.0001).

**Figure 2.**
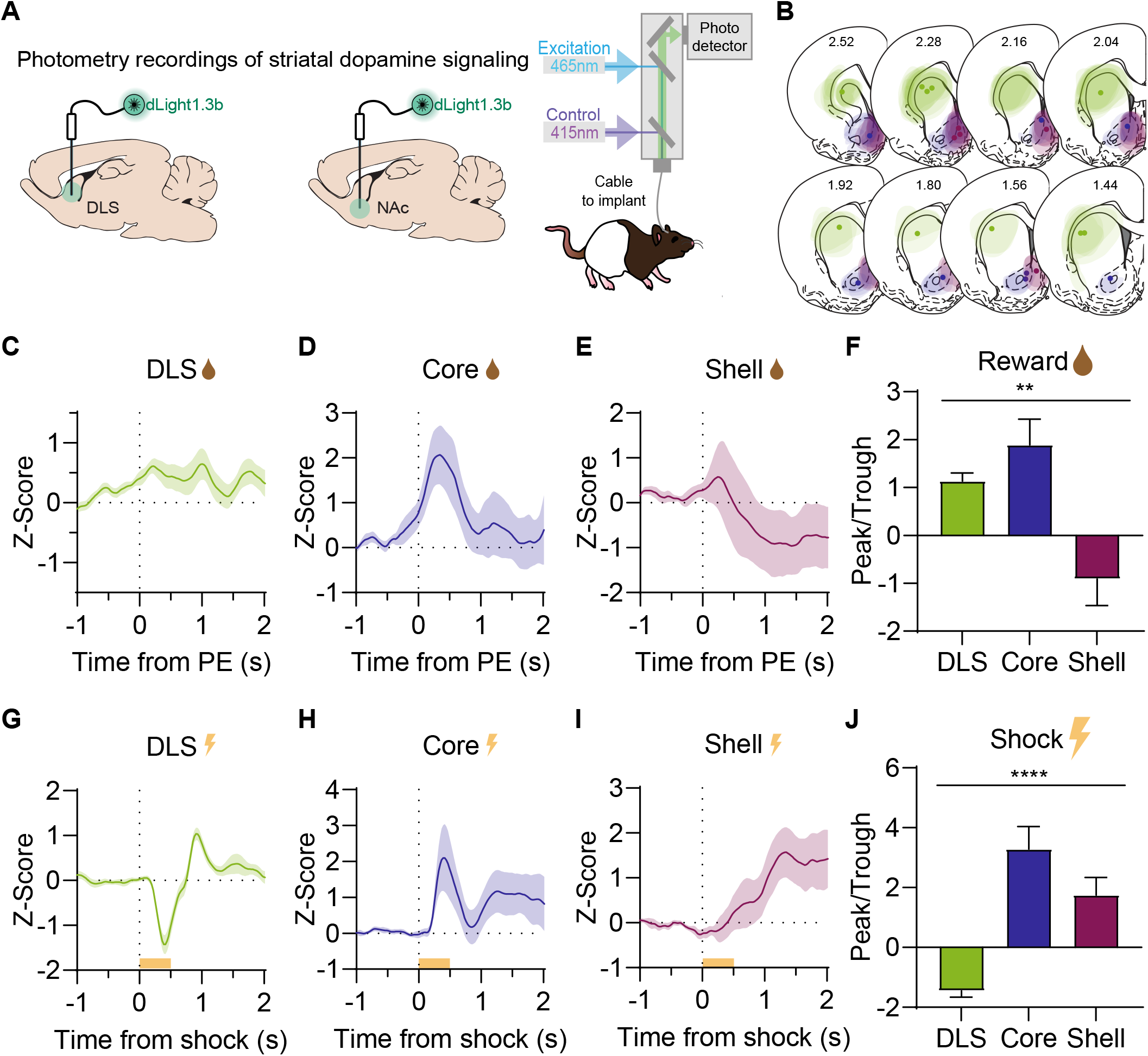
In vivo dopamine recordings across the striatum and responses to oppositely valenced unconditioned stimuli. A The dopamine sensor dLight 1.3b was expressed in the DLS and NAc (shell or core) of wild type Long Evans rats. Schematic of the photometry system; dLight signals were fitted against a control signal to control for photobleaching and movement artifacts. B Photometry measurements were made from three striatal subregions: the DLS (green, n=10), NAc core (blue, n=8), and NAc shell (purple, n=5). C In the DLS, dopamine increased mildly during rewarded port entries (PEs). D In the core, there was a large positive dopamine response during rewarded PEs. E In the shell, there was a slight peak in dopamine followed by a dip below baseline during rewarded PEs. F Quantification of responses plotted in D-F. G In the DLS, dopamine dipped sharply then rebounded above baseline before returning in response to the shock. H In the core, there was a dopamine peak in response to the shock. I In the shell, there was not a clear response to the shock. J Quantification of responses plotted in H-J. All bars represent mean ± SEM. ****p<0.0001, **p<0.01.

### Pavlovian valence ambiguity shapes dynamic heterogeneity in striatal dopamine signals

Dopamine signals in the striatum track elements of appetitive and aversive Pavlovian learning, but many studies examine conditioning in the context of univalent learning conditions. Here, we assessed striatal dopamine under conditions of valence ambiguity, whereby cues that predict positive and negative outcomes were intermingled and rapid discrimination was required for appropriate behavior. We recorded dopamine signals in the DLS, NAc core, and NAc medial shell during the Pavlovian discrimination reversal task described in **Figure 1**.

Focusing first on the CS+R trials (**Figure 3**), heterogeneous dopamine signals emerged across training (**Figure 3B-G**). In the DLS, the initial CS+R response was negative (**Figure 3B**), however, as training progressed, the negative signal diminished (area under the curve [AUC], F(3,27)=8.646, p=0.0003) and became biphasic,with a peak emerging after the initial negative response. In contrast, the DLS response to the reward (**Figure 3C**; first port entry after reward delivery), did not change throughout training (AUC, F(3,25)=0.9476, p=0.4326). In the core, the early response to the CS+R was biphasic, with an initial peak followed by a trough (**Figure 3D**). Across training the trough disappeared and the response became larger (AUC, F(3,21)=12.29, p<0.0001). The core response to reward (**Figure 3E**) did not change throughout training (AUC, F(3,20)=1.307, p=0.2998). The shell response to the CS+R was initially negative, with a gradual progression across training to a large peak (**Figure 3F**, AUC, F(3,12)=15.00, p=0.0002). The shell response to reward (**Figure 3G**) also did not significantly change over the course of training (AUC, F(3,11)=0.5755, p=0.6429). Across all recordings sites, evolution in the shapes of the cue signals were seen comparing sessions 5 to 10/15 (i.e., pre versus post reversal). Together these results highlight subregion-specific dopamine responses to reward-predictive cues, which change as valence discrimination learning progresses.

**Figure 3.**
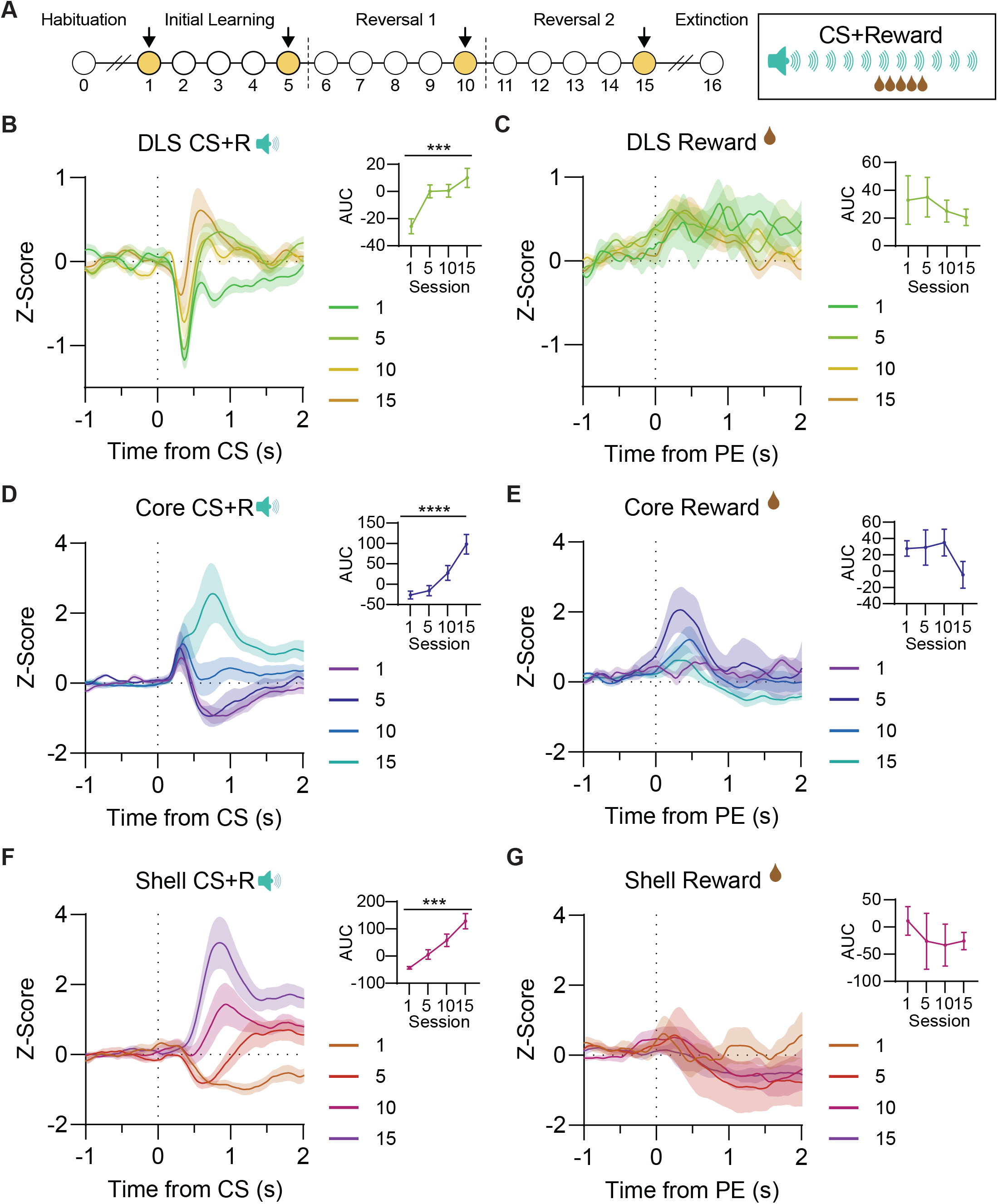
Dynamic, heterogeneous conditioned dopamine signals in response to a reward predictive cue. A Experimental timeline. Photometry data from the highlighted sessions (1, 5, 10, 15) are shown in this figure. Schematic of the CS+R tone showing reward delivery from 10-15 s. In the (B) DLS, (D) core, and (F) shell, dLight responses to the CS+R increased throughout training with different patterns in each striatal subregion. (C, E, G) Reward-associated dLight signals were different in magnitude across subregions but remained stable within each subregion across training. All bars represent mean ± SEM. ****p<0.0001, ***p<0.001.

Beyond the classic connection with reward-related learning and reinforcement, at least some striatal dopamine signals also track aversive learning (Badrinarayan et al., 2012; Lerner et al., 2015; Matsumoto and Hikosaka, 2009; Oleson et al., 2012). To explore this in the context of our valence discrimination task, we next focused on CS+S trials (**Figure 4**), and found heterogeneous dopamine signals to the CS+S and shock across training (**Figure 4B-G**). In the DLS, early in training, the dopamine response to the CS+S was negative (**Figure 4B**). As training progressed there was no consistent change in this signal (AUC, F(3,27)=1.653, p=0.2005). The DLS response to the shock itself had a distinct biphasic shape, with a trough followed by a similar magnitude peak at shock offset that did not change across training (**Figure 4C**; AUC, F(3,27)=1.953, p=0.1448). In the core, the dopamine response to the CS+S showed an initial peak followed by a trough (**Figure 4D**), and this signal also did not significantly change over time (AUC, F(3,21)=2.144, p=0.1251). The core response to the shock itself, which comprised a sharp peak (**Figure 4E**) also did not change (AUC, F(3,21)=0.8604, p=0.4770). The dopamine response to the CS+S in the shell included a rapid decrease with slow recovery to baseline (**Figure 4F**), which trended smaller across training (AUC, F(3,12)=2.730, p=0.0903). While the dopamine response to the shock itself in the shell was relatively small, there was a slow dopamine increase at shock offset (**Figure 4G**) that did not change with training (AUC, F(3,12)=0.2348). Collectively, these results demonstrate a role for subregion-specific dopamine responses to shock-predictive cues that are relatively invariant across valence discrimination learning.

**Figure 4.**
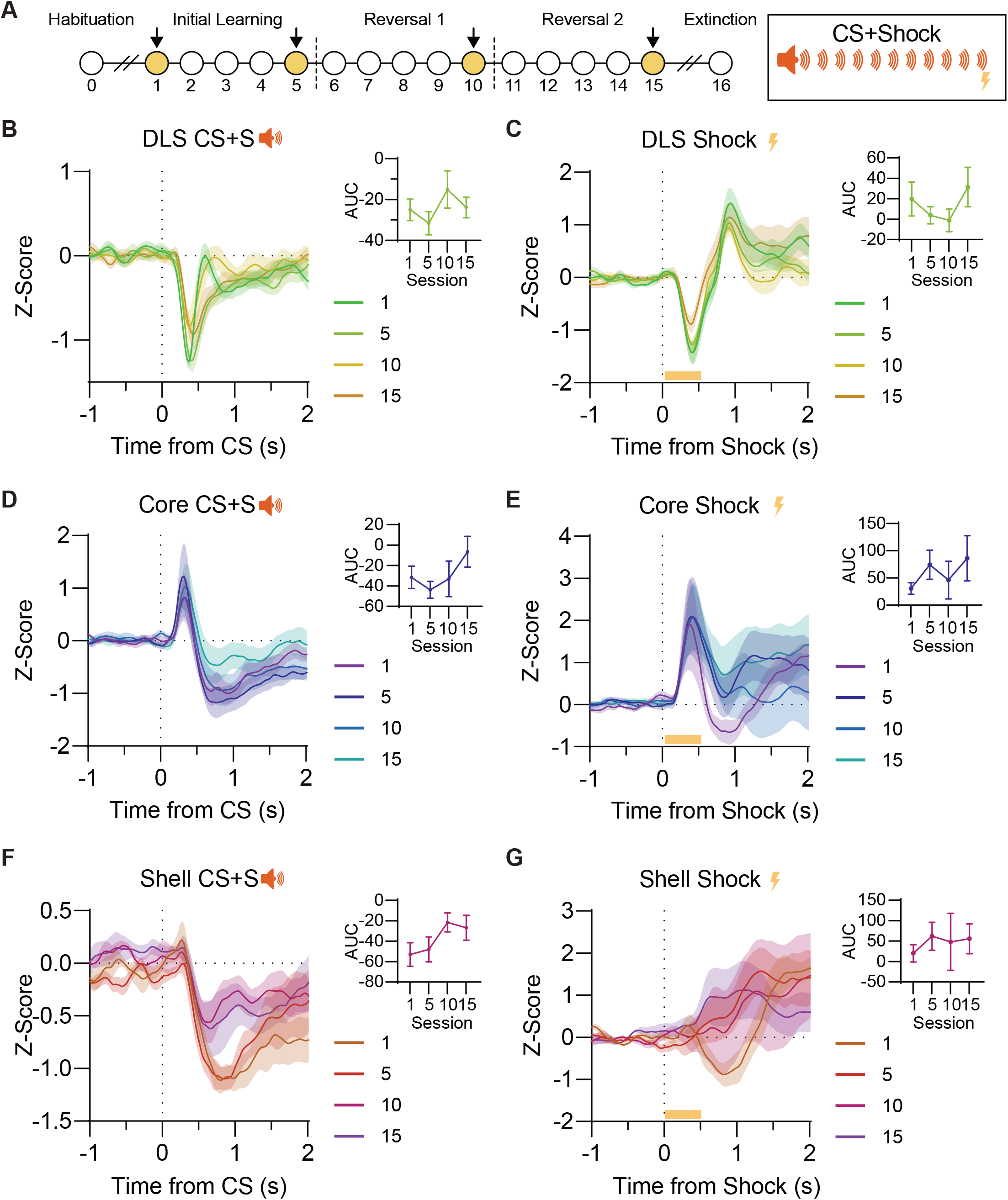
Stable, heterogeneous dopamine signals in response to a threat predictive cue. A Experimental timeline. Photometry data from the highlighted sessions (1, 5, 10, 15) are shown in this figure. Schematic of the CS+S tone showing shock delivery from 19.5-20 s. B In the DLS, the dLight signal in response to the CS+S was negative. C The DLS dLight signal was negative during shock and rebounded to a peak after shock offset. D In the core, the dLight signal in response to the CS+S included a peak followed by a trough. E In the core, the dLight response to the shock was positive. F In the shell, the dLight response to the CS+S was negative. G There was no dopamine response in the shell during shock, but a mild increase at shock offset. All bars represent mean ± SEM.

To compare dopamine signals between reward and shock cues more directly, we calculated a signal bias score based on the ratio of peak (most positive) and trough (most negative) signals measured in the 2 s after onset of either cue type. A positive score reflects a dopamine response where the absolute value of the peak was larger than that of the trough, and a negative score reflects a larger trough than peak. Using this approach, we identified distinct patterns of bias across regions (**Suppl Figure 3**). Across all three regions, initial responses to both cues were biased negative, particularly in the DLS and shell. In the DLS, with training, the reward cue response increased to a neutral position, while the shock cue response remained negative (session x cue type interaction, F(3,27)=5.323, p=0.0052; effect of cue type, F(1,9)=8.216, p=0.019). In the core, the reward cue bias became strongly positive, and the shock cue reached a neutral position (session x cue type interaction, F(3,21)=2.288, p=0.108; effect of cue type, F(1,7)=17.27,p=0.0043). Conversely, in the shell, the reward cue response became strongly positive and the shock cue remained negative across training (session by cue type interaction, F(3,12)=5.125, p=0.0164; effect of cue type, F(1,4)=89.03,p=0.0007).

### Region consistent dopamine responses to the CS-in the multivalent cue discrimination task

Recent evidence suggests that nucleus accumbens dopamine signals track conditioned safety cues, the presence of which signal relief from, or avoidance of, an expected aversive outcome (Luo et al., 2018; Stelly et al., 2019). In our task, the CSpredicted no outcome, but given that it was intermingled with the CS+ cues and was the only cue never paired with shock, within this learning context it likely functioned akin to a safety cue (Day et al., 2016; Ng et al., 2018; Ng and Sangha, 2023). This was behaviorally evident, given that time in port during the CS-was higher than during the CS+S, and freezing during the CS-was lower than both the CS+S and CS+R (**Figure 1**). In contrast to the variable signals seen for each CS+ cues, we found qualitatively similar dopamine responses to the CS-across the striatum (**Suppl. Figure 4B-G**). Early in training, responses to the CS-onset were small and slightly negative. As training progressed, strong positive signals to the CS-emerged across regions (DLS AUC, F(3,27)=7.012, p=0.0012; core AUC, F(3,21)=7.312, p=0.0015; shell AUC, F(3,12)=2.580, p=0.1021). We also saw strong positive dopamine responses in all three regions at the offset of the CS-, which changed in shape but not magnitude across training (DLS AUC, F(3,27)=1.067, p=0.3795; core AUC, F(3,21)=1.796, p=0.1787; shell F(3,12)=1.854, p=0.1913).

### Valence reversal prompts rapid updating of Pavlovian conditioned behavior

In decision making contexts, the meaning of environmental stimuli is often dynamic, and previously learned associations must be updated based on changing contingencies. We probed this in our valence discrimination task by reversing the predictive meaning of reward and shock cues at two points in training (**Figure 5A**). On the first day of reversal 1 (day 6), the tones for the CS+R and CS+S were swapped. On the first day of reversal 2 (day 11), the tones were reversed back to the original contingency from the initial learning phase. Similar to the initial learning phase, there were no sex differences in behavior during the reversal phases (**Suppl. Figure 1**).

**Figure 5.**
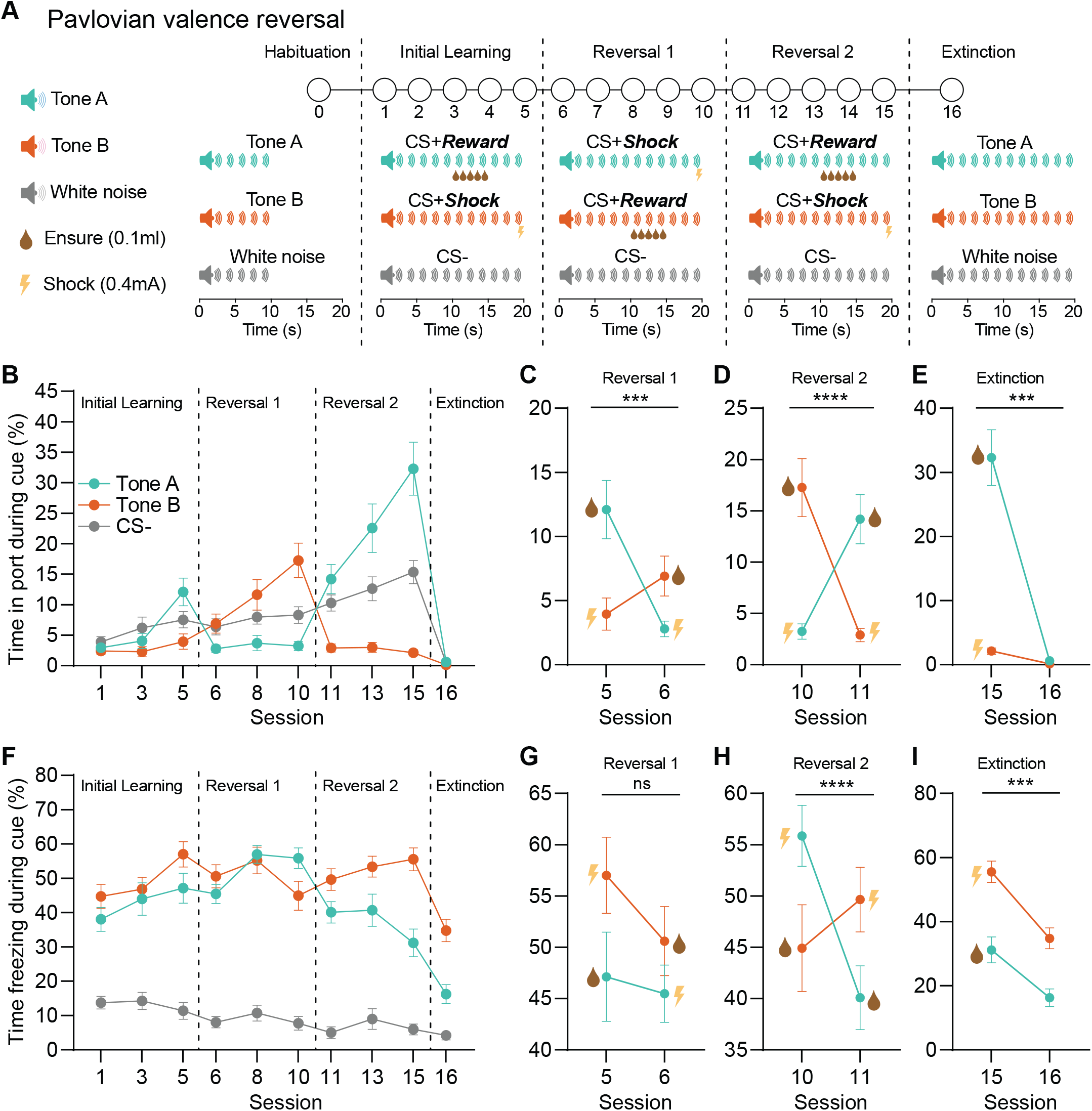
Pavlovian conditioned behavior rapidly updates upon cue reversal. A Experimental timeline. Training began with habituation to the three auditory cues used in the task. Reward (CS+R), shock (CS+S), and neutral cue (CS-) trials were then intermingled, with the CS+R and CS+S switching identity during reversal 1 (days 5-10) before switching back to the original contingency during reversal 2 (days 11-15). The final session (day 16) was conducted under extinction conditions. B Percent time in port during cues diverged throughout training, with behavior updating when tones predicting reward and shock were reversed on sessions 6 and 11, along with extinction. Rats increased time in port when the tone predicted reward, decreased time in port when the tone predicted shock, and remained in the middle when white noise predicted nothing. C Within the first session of reversal 1, rats’ behavior changed due to cue reversal: there was a bias in the reversal, where rats decreased time in port to tone A (reward → shock) more than they increased time in port to tone B (shock → reward). D Within the first session of reversal 2, rats’ behavior rapidly changed back, with an increase in percent time in port from shock → reward reversal and decrease from reward → shock reversal. The bias for reversal from reward > shock that was seen in reversal 1 disappeared. E Percent time in port strongly decreased during extinction. F Percent time freezing during cue diverged throughout training, with behavior appropriately updating when tones predicting reward and shock were reversed on sessions 6 and 11 along with extinction. Rats increased time freezing when the cue predicted shock, decreased time freezing when the tone predicted reward, and remained lowest when white noise predicted nothing. Some generalization to the two tones occurred, as freezing remained higher for reward-predictive cues compared to the CS-. G Within the first session of reversal 1, freezing behavior did not reverse to reflect the new contingencies. H Within the first session after reversal 2, freezing behavior full reversed. Percent time freezing decreased for tone A (shock → reward) more than it increased for tone B (reward → shock). I In extinction, freezing behavior decreased modestly for both cues. All bars represent mean ± SEM. ****p<0.0001, ***p<0.001.

Across training, rats spent more time in the reward port on CS+R trials (**Figure 1, 5B**). In the first session of reversal 1 (day 6), percent time in port updated to reflect the new contingencies (**Figure 5C**; session x tone interaction, F(1,22)=18.80, p=0.0003). The reward → shock transition resulted in a decrease in percent time in port (multiple comparisons test, Tone A Session 5 vs. 6, p=0.0001) but not the shock → reward transition (Tone B Session 5 vs. 6, p=0.1530), indicating that rats were faster at updating port entry behavior when the valence of the cue shifted from positive to negative. By the end of reversal 1, the time spent in port had increased significantly for tone B, reflecting the change in cue valence (multiple comparisons test, Tone B Session 6 vs. 10, p<0.0001), and the time in port remained low for tone A throughout the session (multiple comparisons test, Tone A Session 6 vs. 10, p>0.9999).

On the first day of reversal 2 appetitive behavior rapidly shifted, and rats quickly returned to the pattern of port entry behavior seen in the initial training phase with the original cue contingencies (**Figure 5D**; day 10 vs day 11, session x tone interaction, F(1,22)=33.37, p<0.0001). The difference in reversal speed between the CS+R and CS+S disappeared, as behavior updated to both Tone A (multiple comparisons test, p=0.0019) and Tone B (p=0.0001) within this first session. Following the retraining phase after reversal 2, we conducted a final session under extinction conditions (day 16), wherein the CS+R and CS+S were presented without outcomes. During extinction, time spent in the port dropped dramatically (**Figure 5E**; session x tone interaction, F(1,22)=51.44, p<0.0001). Overall, time spent in the port showed a tight coupling with the CS+ cue contingencies across sessions, selectively increasing in response to the reward-paired cue and rapidly decreasing to the same cue when it was paired with shock.

Across training, conditioned freezing was generally more prevalent on CS+S trials (**Figure 1, 5F**). However, we found that freezing behavior was slower to update than port entry behavior upon cue reversal. On the first session of reversal 1, freezing to either CS+ did not change, despite the shift in contingencies. Freezing discrimination to the reversed cues did eventually emerge by the end of training after reversal 1 (**Figure 5 F,G**; day 10), and subsequently, freezing adapted quickly on the first session of reversal 2 when cue contingencies returned to their original status (**Figure 5H**; day 10 vs day 11, session x tone interaction, F(1,22)=23.21, p<0.0001). During extinction, freezing decreased to both cues (**Figure 5I**; two-way ANOVA, session main effect, F(1,22)=84.23, p<0.0001), but remained higher for the CS+S than the CS+R (cue type main effect, F(1,22)=79.15, p<0.0001). Together, our results indicate that Pavlovian cue conditioned behavior is generally flexible upon changing valence predictions, but threat-related responding is slower to update than reversal of reward-related responding.

### Subregion-specific updating of striatal dopamine signals after valence reversal

We assessed how striatal dopamine responses to the CS+R and CS+S changed when the meaning of the cues was reversed (**Figure 6A**). On the last day of initial training (session 5), the dopaminergic response in the DLS to both the CS+R and CS+S were characterized by sharp troughs. This was followed by a small peak to the CS+R while the CS+S response was followed by a slow return to baseline (**Figure 6B**), resulting in an overall larger decrease in dopamine in response to the CS+S than the CS+R in the DLS (AUC, paired t-test, t(9)=7.196, p<0.0001). On the first day of reversal (day 6) the response profiles in the DLS were similar to day 5: the tone originally predicting shock continued to evoke a stronger decrease in dopamine, despite now predicting reward (**Figure 6C**; AUC paired t-test, t(9) = 2.971, p=0.0157; **Figure 6D**; two-way ANOVA, tone main effect, F(1,9)=30.24, p=0.0004). Thus, upon reversal the DLS responses to each tone remained constant despite their change in predictive nature.

**Figure 6.**
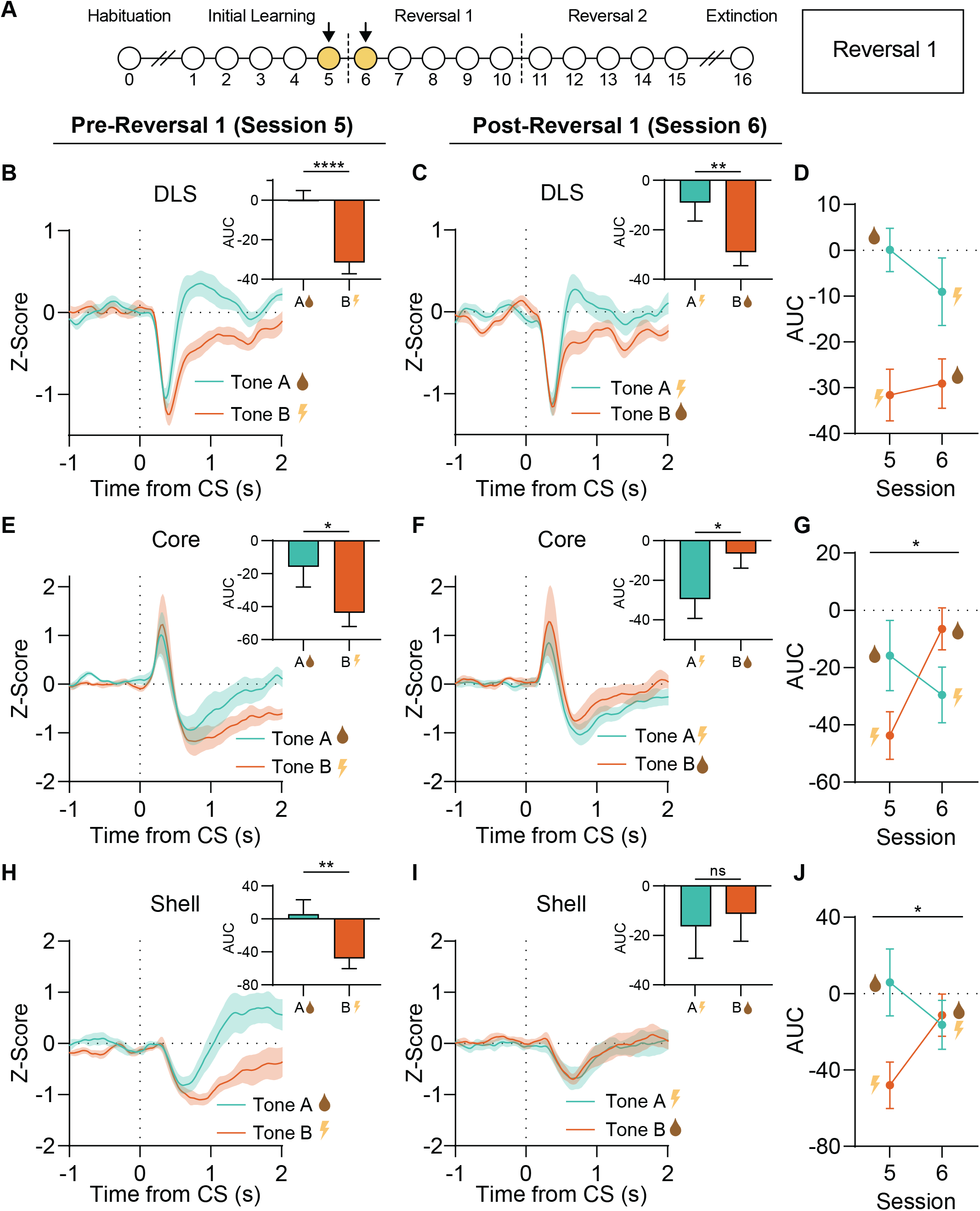
Subregional striatal dopamine signals update to valence reversal at different rates. A Experimental timeline with the highlighted sessions outlined in this figure. B The dopamine signals in the DLS in response to tone A (CS+R) and tone B (CS+S) differed by session 5. The response to the CS+R included a trough followed by a shallow peak, while the response to the CS+S included a trough with a slow return to baseline; the difference is quantified as the area under the curve (AUC). C Upon reversal 1, the dopamine signal in response to tone A (now CS+S) still includes a trough followed by a soft peak while the response to tone B (now CS+R) still includes a trough with a slow return to baseline. The AUC for tone A remains higher than that for tone B, even though the tones have reversed in meaning. D The DLS did not show an updated response only one session after reversal, because the AUC in response to tone A (CS+R → CS+S) remained higher than that to tone B. (CS+S → CS+R). E During session 5 the dopamine signals in the core in response to both tone A and B included a peak followed by a trough, but the response to tone A included a faster return to baseline. F With reversal 1, the dopamine signals in the core in response to tones A and B again included a peak followed by a trough, but now the response to tone B includes a shallow trough with a faster return to baseline. G The core showed updated dopamine responses to tones A and B within the first session of reversal, with the AUC increasing from CS+S → CS+R and decreasing form CS+R → CS+S. H During session 5 the dopamine response in the shell to tone A (CS+R) included a soft trough followed by a soft peak, while the response to tone B (CS+S) included a soft trough with a slow return to baseline. I Upon reversal in the shell, the dopamine responses to tones A and B include a soft trough with a return to baseline. J The shell did not display a fully updated response to the cue reversal, but it began to change. The response to tone A (CS+R → CS+S) decreased while the response to tone B (CS+S → CS+R) increased. All bars represent mean ± SEM. ****p<0.0001, ***p<0.001, *p<0.05.

During session 5, the response in the core to both the CS+R and CS+S included a sharp peak followed by a small trough and a return to baseline, but the return to baseline was slower in response to the CS+S (**Figure 6E**). Similar to the DLS, the dopamine response in the core to the CS+S was larger than for the CS+R (AUC paired t-test, t(7) = 2.414, p=0.0465). However, unlike the DLS, the dopamine response in the core after reversal to the new CS+R was significantly reduced compared to the new CS+S (**Figure 6F**; AUC paired t-test, t(7) = 3.302, p=0.0131). Comparing sessions 5 and 6 for the core, cue-evoked signals updated during the first day of reversal 1 (**Figure 6G**; session x tone interaction, F(1,7)=11.97, p=0.0106), which was driven by an increased response to Tone B (p=0.0090), but not Tone A (p=0.2292).

In the shell on session 5, the response to the CS+R included a delayed trough followed by a slow positive response, while the CS+S induced a larger trough that was delayed a greater extent (trough, paired t-test, t(4)=5.280, p=0.0062; trough latency, t(4)=3.405, p=0.0272) followed by a slow return to baseline (**Figure 6H**). As in the DLS and core, the dopamine response in the shell was larger to the CS+S than the CS+R (AUC, paired t-test, t(4)=7.096,p=0.0021). Upon reversal, the response in the shell to both the CS+R and CS+S became similar (**Figure 6I**; AUC paired t-test, t(4)=0.4480, p=0.6774). However, comparing signals across sessions 5 and 6 revealed that both signals changed, becoming more similar (**Figure 6J**; interaction, F(1,4)=20.04, p=0.0110). Thus, while shell dopamine did not fully reverse by session 6, it was in the process of updating, unlike the DLS. Multiple comparisons of the AUC for Tone A did not significantly change between sessions 5 and 6 (p=0.0752), but the AUC increased for Tone B (p=0.0169).

### Serial cue reversal effects on valence encoding across dopamine subregions

Rats were trained on the new cue contingencies for 4 more sessions (**Figure 7A**). In the session before reversal 2 (session 10), the dopaminergic response to the CS+R and CS+S in the DLS included a rapid decrease in dopamine with a quick return to baseline (**Figure 7B**). Despite this additional training, the DLS cue responses never fully updated its response to the changing contingencies upon reversal 1 (AUC, paired t-test, t(9)=1.945, p=0.0836), although a trend in cue discrimination had emerged by session 10 (**Figure 7B**). On reversal 2 (session 11, cue contingencies returned to their original identity), the DLS response to the CS+R included a sharp trough followed by a sharp positive peak, while the response to the CS+S only included a sharp trough (**Figure 7C**). Although there was only a trend toward cue discrimination on session 11 (**Figure 7C**; AUC; paired t-test, t(9)=1.907, p=0.0889), comparing sessions 10 and 11 showed a change in the dopamine signal as a function of session and tone upon reversal 2 (**Figure 7D**; session x tone interaction, F(1,9)=8.363, p=0.0178), indicating the response in the DLS was in the process of updating/reversing. By session 10, the dopamine signal in the core to the CS+R was larger than the CS+S (**Figure 7E**; AUC, paired t-test, t(7)=3.079, p=0.0178). Upon reversal 2 (session 11), these cue signals became more similar (**Figure 7F**; AUC, paired t-test, t(7)=1.056, p=0.3259), but comparison of session 10 and 11 suggested that the core signal was in the process of updating/reversing (**Figure 7G**; session x tone interaction, F(1,7)=8.860, p=0.0206). During session 10, the dopamine response in the shell to the CS+R was larger than for the CS+S (**Figure 7H**; AUC paired t-test, t(4)=3.841, p=0.0184). Upon reversal (session 11), these signals were qualitatively updated to reflect the renewed contingencies. As with the DLS and core, the shell signal did not fully discriminate cue type on session 11 (**Figure 7I**; AUC paired t-test, t(4)=2.047, p=0.1101), but comparison of sessions 10 and 11 again suggested that the shell signal was in the process of updating (**Figure 7J**; session x tone interaction, F(1,4)=9.581, p=0.0364).

**Figure 7.**
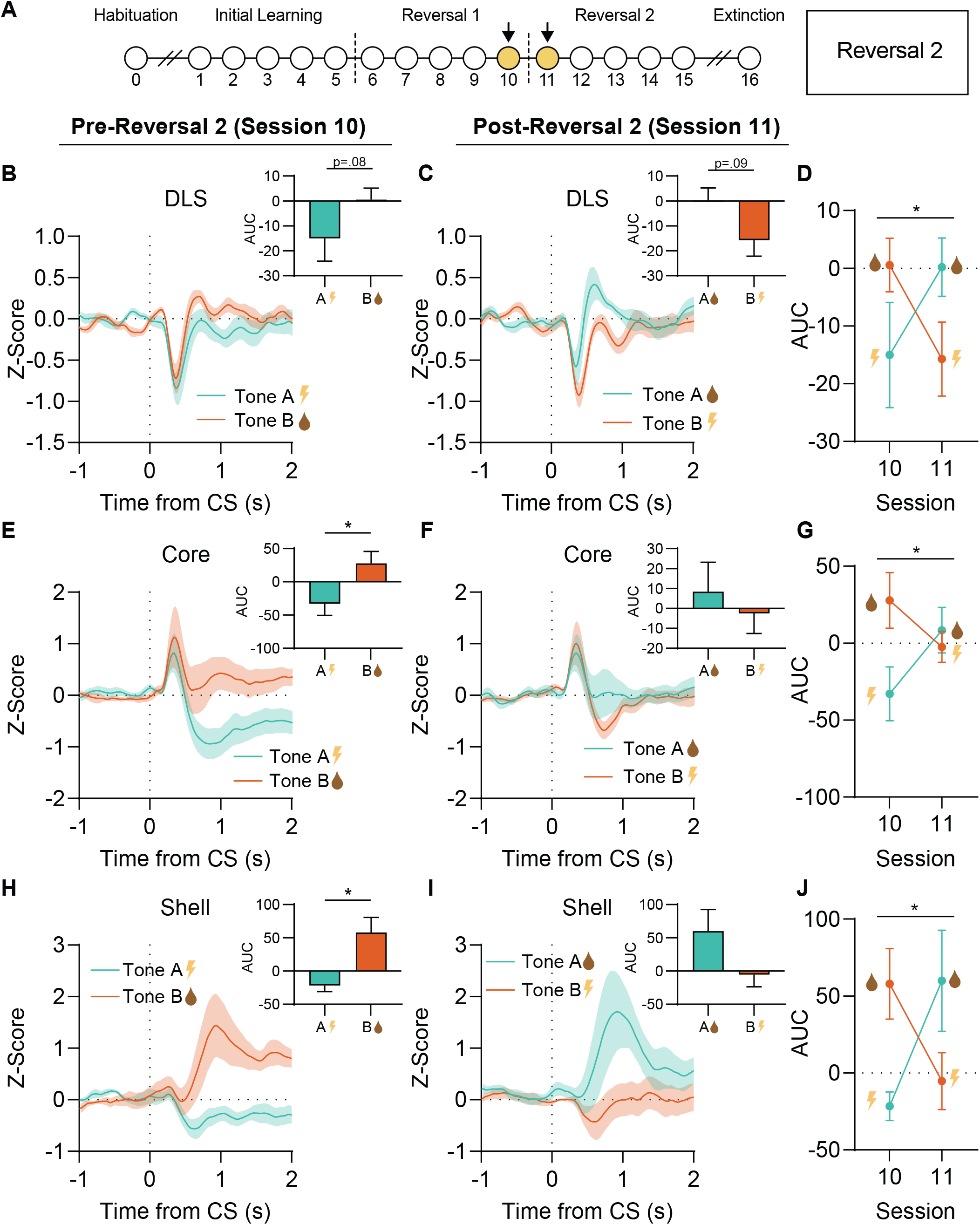
Serial cue reversal effects on dopamine signal updating across striatal subregions. A Experimental timeline highlighting the sessions outlined in this figure. B In the DLS on session 10, the dopamine response to tone A (CS+S) included a trough with a return to baseline while the response to tone B (CS+R) included a trough followed by a peak. C After the second reversal, the cues returned to their original meanings and the dopamine response in the DLS to tone A (CS+R) included a trough followed by a peak while the response to tone B (CS+S) only included a trough. D Comparing the patterns of responding to each cue on days 10 and 11, the DLS signals updated to reflect the new contingencies. E The dopamine response in the core on session 10 to tone A (CS+S) included a peak with a dip below baseline while the response to tone B (CS+R) included a peak with a return to baseline. F Upon reversal 2 in the core, the dopamine signal in response to tone A (CS+R) included a peak. with a return to baseline, while the response to tone B (CS+S) included a peak with a dip below baseline. G Comparing the patterns of responding to each cue on days 10 and 11, the core signals were in the process of updating to the new contingencies. H In the shell on session 10, the dopamine signal in response to tone A (CS+S) included a trough, while the response to tone B (CS+R) included a large peak. I Upon reversal 2 in the shell, tone A (CS+R) developed a large peak while tone B (CS+S) developed a negative response. J Comparing the patterns of responding to each cue on days 10 and 11, the shell signals updated to reflect the new contingencies. All bars represent mean ± SEM. *p<0.05.

### Different rates of dopamine extinction to reward versus threat cues across striatal subregions

Following the second reversal, rats were trained with the original cue associations for an additional 4 sessions (**Figure 8A**). In general, this led to stronger cue discrimination in dopamine signals across all regions in the final session before extinction (session 15). Dopamine signals in response to the CS+R and CS+S were clearly distinct in all regions (**Figure 8B, 8E, 8H**; DLS AUC paired t-test, t(9)= 3.429, p=0.0075; core AUC paired t-test, t(7)=3.865, p=0.0062; shell AUC paired t-test, t(4)= 6.751, p=0.0025).

**Figure 8.**
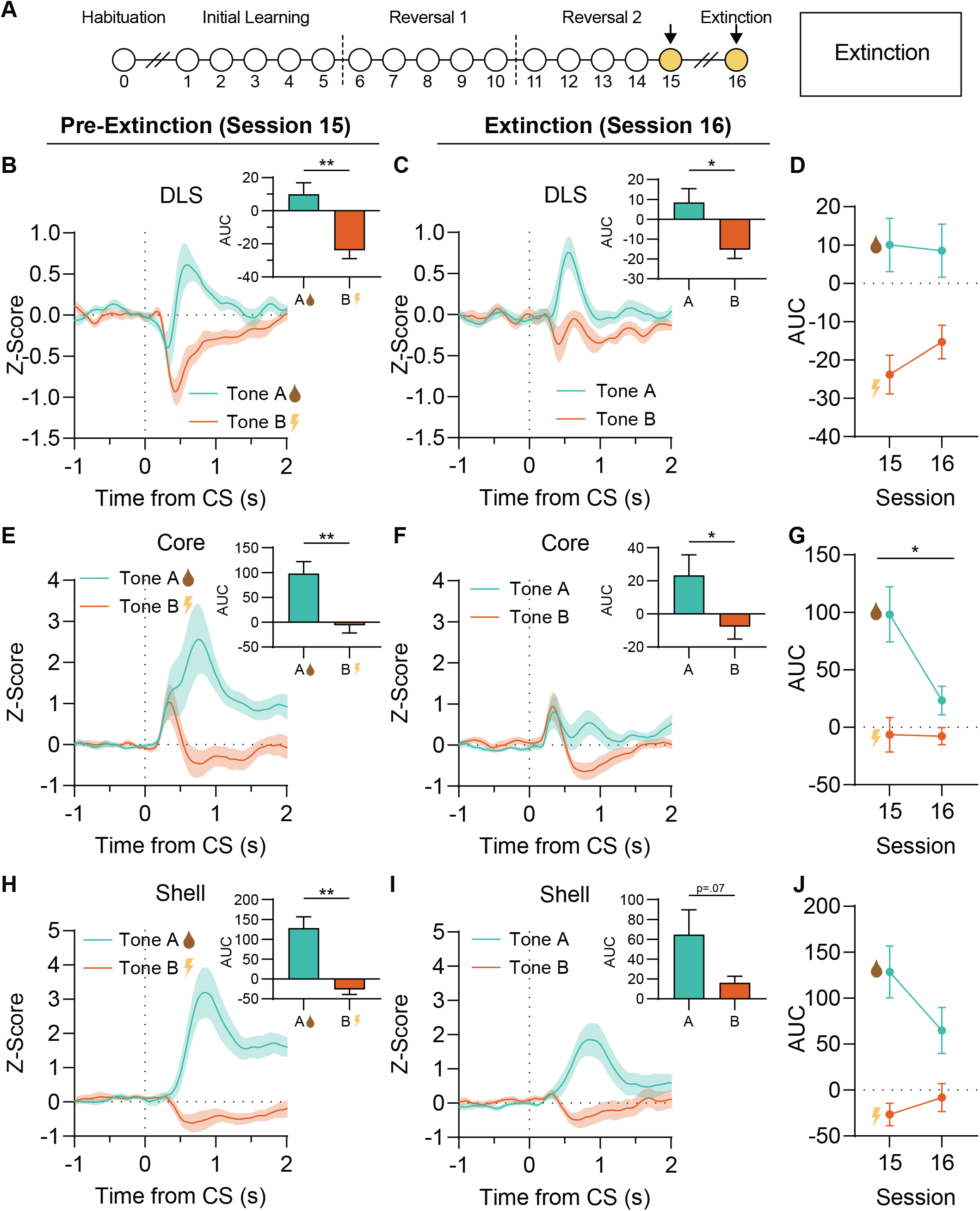
Variable extinction of striatal dopamine responses to reward versus threat cues across striatal subregions. A Experimental timeline showing the sessions highlighted in this figure. B By the final day of training in the DLS, the dopamine signal in response to the CS+R included a trough followed by a peak, while the response to the CS+S included a trough with a slow return to baseline. C After extinction in the DLS, the dopamine signal in response to tone A (previously CS+R) only included a peak. The response to tone B (previously CS+S) became less distinct, with the trough disappearing. D Comparing the last day of conditioning (15) with extinction (16), the DLS signals to either cue did not significantly change. E In the core during the last training session, the dopamine signal in response to tone A (CS+R) included a large peak while the response to tone B (CS+S) included a small peak. F After extinction in the core, the dopamine peak to the previously rewarded cue. was much smaller. G Comparing days 15 and 16, only the reward cue response diminished with extinction. H On the final session of training in the shell, the dopamine signal in response to tone A (CS+R) included a large peak, while the response to tone B (CS+S) included a small dip below baseline. I After extinction in the shell, the dopamine response to the previously rewarded cue was smaller. J Comparing days 15 and 16, shell dopamine signals only modestly extinguished. All bars represent mean ± SEM. **p<0.01, *p<0.05.

During extinction (session 16), DLS dopamine signals in response to the CS+R remained stable (**Figure 8D**; AUC, session 15 vs 16, multiple comparisons test, p=0.7553), while the response to the CS+S became smaller (**Figure 8D**; trough, paired t-test, session 15 vs 16, t(9)=2.351, p=0.0432). DLS dopamine continued to discriminate cue types during extinction (**Figure 8C**; paired t-test, t(9)= 2.565, p=0.0304) and when comparing across sessions 15 and 16 (**Figure 8D-8E**; no interaction, significant main effect of tone, F(1,9)=10.28, p=0.0107).

In the core during extinction, the CS+R signal shrank but the CS+S signal remained stable (**Figure 8F**). The core signals in response to the CS-R showed the most change in extinction, and a comparison of sessions 15 and 16, revealed that responses varied as a function of both session and tone (**Figure 8F-8G**; session x tone interaction, F(1,7)=9.991, p=0.0056). Despite these changes, the cue signals remained distinct after extinction (AUC; paired t-test, t(7) = 2.706, p=0.0304).

Finally, during extinction, the shell response to the CS+R decreased (AUC, multiple comparisons test, trend, p=0.0615) while the CS+S signal remained unchanged (AUC, multiple comparisons test, p=0.4966). This resulted in marginal discrimination in the dopamine signal between cue types (paired t-test, t(4)=2.452, p=0.0703). Comparing sessions 15 and 16 in the shell, the results were similar to the DLS, where dopamine signal in response to the CS+R remained higher than to the CS+S (**Figure 8J**; tone main effect, F(1,4)=32.35, p=0.0047).

## DISCUSSION

We investigated behavioral flexibility and dopaminergic encoding in a task where positive and negative Pavlovian associations were intermingled and serially reversed. From the current data, we highlight four primary conclusions. First, rats readily learned to distinguish between cue types, but reversal sessions suggested a bias in the speed of updating for reward vs threat associations. Second, by recording dopamine signals across three major striatal subregions (DLS, NAc core, and NAc medial shell), we show substantial heterogeneity in responses of reward and threat associations, bolstering the notion that striatal niches signal different aspects of learning and stimulus meaning. While dopamine in all regions discriminated cue types, the direction, timing, and qualitative pattern of that signal was different in each area. Third, our Pavlovian learning context, in which cues predicting positive and negative outcomes are intermingled, promoted unique, multi-phasic dopamine signaling patterns, compared to previous studies involving single valence learning contexts. Ambiguity in perceived cue identity was apparent across all striatal regions, especially early in training, where most dopamine signatures following cue onset included both a negative and positive inflection point. Fourth, we show that some conditioned dopamine signals are quite dynamic to changes in expected valence: most responses updated within a single session of cue reversal, an adaptation that was even more robust upon a second reversal. Together, our results suggest that striatal dopamine encodes multi-valent learning contexts, offering insight into a number of ongoing questions regarding dopamine’s role in signaling different facets of learning, as well as the broad heterogeneity and dynamic nature of the striatal dopamine system.

Notably, our data showed that dopamine in each region encoded valence, as the signal to the reward cue was uniformly more positive than the signal in response to the shock cue. This is broadly consistent with classic notions of dopamine’s role in reward and reinforcement (Schultz et al., 1997; Wise, 2004), as well as more recent investigations into dopamine subregions (van Elzelingen et al., 2022). Recent work has begun to broaden that framework, however, suggesting that at least within the NAc core, dopamine signals salience rather than valence (Bromberg-Martin et al., 2010; Ventura et al., 2007), serving to integrate strength, stimulus intensity, and novelty (Kutlu et al., 2021). Our results support this framework, given that we found strong positive dopamine responses in the core within the first ~1 second after cue and stimulus onset in both reward and shock conditions, throughout training. This suggests that increased core dopamine denotes the presence of an important stimulus (Horvitz, 2000; Redgrave et al., 1999; Ungless, 2004) rather than providing information regarding its valence per se. Similar to other regions, signals in the NAc core readily discriminated between reward and shock cues, however, which was evident in examination of the later part of the response signal (~1-2 seconds after cue onset). For shock cues, this second phase of the core signal reflected a decrease in dopamine that was maintained across training. In contrast, decreases in dopamine in response to the reward cue disappeared as training progressed. Thus, the core dopamine signal likely reflects some integration of salience and valence, potentially transmitted on different timescales early in the cue period, as ambiguity over the stimulus identity is resolved. We saw evidence of distinct biphasic dopamine signals in the shell and DLS as well, although the directionality of dopamine changes in those regions in response to the CS+R and CS+S more closely resembled their valences at the end of training a net positive increase in dopamine in response to the CS+R and a net negative decrease in dopamine in response to the CS+S. This pattern in the shell and DLS suggests the prioritization of cue value/valence over salience. Our data collectively indicate that functional heterogeneity in dopamine signaling exists not just across different anatomical regions, but also in brief temporal windows within a given region.

Our studies underscore the importance of considering more complex decision making contexts involving multiple learned associations, in this case where motivation must be derived from stimulus ambiguity, which is variable trial to trial. A multivalent, dynamic learning landscape presents more opportunities for error signals, as more sensory elements in the environment carry information, and dopamine under these conditions may represent an integration of multiple factors, including errors in predictions of identity (Keiflin et al., 2019; Sharpe et al., 2017; Takahashi et al., 2017). A relatively unique feature of our data was that across recording sites, cue onset usually resulted in a multiphasic response, where the dopamine signal included an increase above and dip below baseline within the first couple seconds. This pattern suggests that stimulus identity was ambiguous, resulting in both CS types evoking elements of reward and threat-related representations simultaneously. As conditioning progressed, cue signals generally became less multi-phasic, suggesting that their predictive identity became more rapidly discernable with learning. Notably, while all recording sites showed some degree of biphasic dopamine responses, the order of negative versus positive deflections differed, which suggests an inherent bias in the representation of valence across striatal dopamine niches.

The resolution of stimulus uncertainty is itself rewarding, and the basal ganglia is centrally implicated in this process (Bromberg-Martin and Monosov, 2020; Gottlieb et al., 2014; Vellani et al., 2020). The offset/end of cues in our task was especially meaningful, because they resolved any remaining ambiguity regarding cue identity. We found cue offset signals in a number of conditions in our data, including in response to the termination of the CS-. Cue offset signals have been seen in other conditions, including at offset of a rewarding cue (Kalmbach et al., 2022) and at shock omission (Kutlu et al., 2023). Cue offset dopamine may also reflect a “relief” or safety signal in our task (Luo et al., 2018; Oleson et al., 2012; Stelly et al., 2019), either confirming that shock will not occur, or that its occurrence is over. In task structures involving uncertainty, dopamine’s behavioral impact may reflect the accumulation of evidence that helps produce the appropriate behavioral response (Beste et al., 2018; Fraser et al., 2023; Lak et al., 2020, 2017; Mikhael et al., 2022). Overall, this complexity underscores the difficulty in assigning specific meaning to a given dopamine signal, because it likely simultaneously represents a multifaceted state incorporating value, salience, novelty, evidence, and other factors.

Recent work suggests that accumbens core dopamine plays a key role in resolving cue-based uncertainty regarding reward prediction (Fraser et al., 2023; Mikhael et al., 2022). Our results add to this notion to suggest that rapid fluctuations in dopamine across striatal regions tracks stimulus identity and valence on a trial-by-trial basis, which could aid in the disambiguation of the appropriate motivational state in that moment. While dopamine in all regions discriminated reward and shock cues, updating of these signals upon cue reversal was faster for the nucleus accumbens recording sites, compared to the DLS, suggesting that DLS dopamine may track a relatively longer running rate of conditioned cue value that is less tied to moment-to-moment stimulus changes (Fiorillo et al., 2003; Hart et al., 2015; Kim et al., 2015). Consistent with this, DLS dopamine signals showed the least change during extinction in our study. The relative slowness of DLS updating is perhaps consistent with classic notions of the role of this region in stimulus-response, habit-like or inflexible behavior (Everitt and Robbins, 2005; Hart et al., 2015; Klanker et al., 2017; Lerner, 2020; Malvaez and Wassum, 2018; Nestler and Lüscher, 2019; Yin et al., 2009; Yin and Knowlton, 2006). Interestingly though, we saw robust extinction of behavioral responses, and so persistent cue-evoked DLS dopamine signals in this case may partly reflect local constraints on signaling dynamics, rather than a neural readout of habit per se.

Our results support the conclusion that the striatum represents valence, with broad heterogeneity. This is generally consistent with previous studies using a variety of tasks and recording methods, but we find some discrepancies. For example, previous studies have shown that dopamine neurons projecting to DLS are excited by aversive stimuli, while dopamine neurons projecting to the ventral striatum are inhibited by aversive stimuli (Lerner et al., 2015; Matsumoto and Hikosaka, 2009). In other studies, dopamine terminals and dopamine release in the medial shell are excited by aversive stimuli, while inhibited in other NAc subregions (Badrinarayan et al., 2012; De Jong et al., 2019; Goedhoop et al., 2022; Oleson et al., 2012). Yet other studies have reported uniformly negative dopamine signals across striatum in response to aversive stimuli (van Elzelingen et al., 2022). We found a different set of response patterns here, where dopamine in the DLS showed a biphasic, negative-then-positive response, the core showed a positive response, and the shell showed no clear response to shock. In a discriminative stimulus task using fast-scan cyclic voltammetry, the core, not the shell or DLS, produced phasic positive dopamine signals in response to a reward-predictive stimulus (Brown et al., 2011). In contrast, here we found that the DLS, core, and shell developed positive responses to a reward-predictive stimulus, although it took longer for this signal to emerge in the DLS. A number of important methodological differences exist across these studies, and more work is required to further understand how different striatal regions signal valence, but our results suggest that multivalent, conflict-based, or valence-ambiguous learning contexts (Burgos-Robles et al., 2017; Kutlu et al., 2020) may produce unique dopamine response patterns compared to other learning environments.

Here, by measuring striatal dopamine signals and behavior during a multivalent Pavlovian discrimination task, we uncover region-specific patterns of dopamine signaling. We show that the DLS, core, and shell encode cues of positive and negative valence with differing response profiles, and valence ambiguity dynamically reshapes these dopamine signals as contingencies change. This work provides new insight into the striatal involvement in learning and value-based decision making.

## Acknowledgements

This work was supported by NIH grants T32 MH115886 (TJO) and R00 DA042895, R01 MH129370, and R01 MH129320 (BTS). We thank all members of the Saunders lab for helpful discussion surrounding this project.

## Author Contributions

Design and conceptualization: KNB, TJO, BTS. Data collection: KNB, JP, TJO. Data analysis and visualization: KNB, JP, TJO, ARW, BTS. Manuscript preparation: KNB, ARW, BTS.

## METHODS

### Subjects

Outbred female (n = 11; ~8 weeks old, 250-274 g at arrival) and male (n = 12; ~8 weeks old, 300-324 g at arrival) Long-Evans rats (Envigo; Indianapolis, Indiana, USA) were single-housed in a humidity- and temperature-controlled vivarium, with ad libitum access to water. Rats were acclimated to the vivarium for at least five days and handled for at least three days prior to any procedures. Rats were food-restricted to ~90% of free-feeding weight one week prior to starting any behavioral experiments. All procedures were done in accordance with the National Institutes of Health’s Office of Laboratory Animal Welfare and the University of Minnesota’s Institutional Animal Care and Use Committee.

### Stereotaxic surgery

Rats were anesthetized with isoflurane (5% induction, 1-2% maintenance; Patterson Veterinary) and injected with carprofen (5 mg/kg, 50 mg/ml, sc), cefazolin (70 mg/kg, 0.2 g/ml, sc), and saline (0.5 ml, sc). Following head-fixation in a digital stereotax (Kopf Instruments), the skull was exposed, leveled, and scored, and a craniotomy was drilled over the target region. Plasmid containing dLight viral DNA (pAAV-CAG-dLight1.3b) was purchased from Addgene (#125560; Patriarchi et al., 2018) and packaged by the University of Minnesota’s Viral Vector and Cloning Core with a titer of ~1x1013 GC/ml (AAV5-CAG-dLight1.3b). Viral vectors were loaded into 10μl gas-tight syringes (Hamilton Company) and infused (700 nl, 140 nl/min) unilaterally into the nucleus accumbens medial shell (coordinates in mm, relative to Bregma: AP +1.5, ML −1.0, DV −7.5), nucleus accumbens core (AP +1.3, ML −1.3, DV −7.0) or dorsolateral striatum (AP +1.0/1.2, ML −3.3/3.5, DV −5.0). Mono fiber optic cannulae (silica/polymer, 400 μm, NA 0.48, 9.0mm fiber, 2.5 mm metal ferrule, flat tip; Doric Lenses Inc., Quebec, Canada) were implanted unilaterally into the nucleus accumbens medial shell (AP +1.5, ML −1.0, DV −7.3), nucleus accumbens core (AP +1.3, ML −1.3, DV −6.8), or dorsolateral striatum (AP +1.0/1.2, ML −3.3/3.5, DV −4.8) above injection sites. Fiberoptic cannulae were secured with dental cement (Lang Dental; Wheeling, Illinois) and secured with self-tapping bone screws (Fine Science Tools; Foster City, California) on the skull. Following surgery, rats were given daily injections of carprofen (5 mg/ kg, 50 mg/ml, sc) and cefazolin (70 mg/kg, 0.2 g/ml, sc) for three days.

### Habituation and Pavlovian Conditioning

Behavior training began approximately 3-4 weeks after surgery. Chambers (Med Associates) were outfitted with speakers for the delivery of tone (high: 4500 Hz, low: 2900 Hz) or white noise cues, a syringe pump to deliver liquid reward, a magazine port equipped with an infrared beam to sense port entries, and floor bars for the delivery of electrical shocks. Chambers were cleaned with 70% ethanol and bedding was replaced after each session.

Rats were first acclimated to the behavioral chambers, conditioning cues, and optic cable tethering in a ~35-min habituation session. During this session, rats were tethered to a cable and given 15 noncontingent cue presentations (~60dB; 5 high tone, 5 low tone, 5 white noise) on a 120-s average variable time (VT) schedule (range 90-150 s) lasting 5-s each. These cue presentations were not associated with any outcome.

Next followed a Pavlovian reversal task adapted from previous studies (Burgos-Robles et al., 2017). During the initial learning phase, (five training sessions), rats were presented with 60 cues on a 60 s average VT schedule (range 45-75 s), with each cue presentation lasting 20 sec. 25 cues predicted liquid reward (CS+R, 0.1 mL Ensure chocolate shake) delivered from 10-15 s, 25 cues predicted footshock (CS+S, 0.4 mA) delivery from 19.5-20 s, and 10 cues had no outcome (CS). Cues were presented in a pseudorandom order. The high and low tones were counterbalanced across rats for the CS+R and CS+S, and white noise was always used for the CS-. Each session lasted ~80 min, and port entry instances and the duration of each port entry throughout each session were recorded by Med Associates software. Cameras positioned above each chamber recorded movement throughout each session for behavioral analysis.

### Cue-outcome Reversals

After five sessions of initial learning, the reward and shock cue contingencies were reversed (reversal 1), whereby the CS+R became the CS+S and vice versa (e.g. a tone that predicted reward now predicts shock). All other aspects of the task were identical to the initial learning phase. After five sessions in the reversal 1 phase, the contingencies were reversed again back to the original associations (reversal 2) as in the initial learning phase for another five sessions.

### Extinction

Behavioral testing concluded with one extinction session. This session was identical to the Pavlovian reversal task, except with the omission of the reward and shock deliveries.

### Fiber photometry recordings

To measure dopamine signals in the core, shell, and DLS, fiber photometry was performed using a system with optical components from Doric Lenses (Québec, Canada) controlled by a real-time processor from Tucker-Davis Technologies (RZ5P, TDT, Alachua, FL) running Synapse software for session control and data acquisition, similar to previous studies (Saunders et al., 2018). A fluorescence mini-cube transmitted light streams from 465 nm and 415 nm LEDs, sinusoidally modulated at 211 Hz, and 330 Hz, respectively. LED currents were adjusted to ~50 μW for each signal. Fluorescence from neurons at the fiber tip was transmitted back to the minicube, where it was passed through a GFP emission filter, amplified, and focused onto a high sensitivity photoreceiver (Newport, Model 2151). The RZ5P processor running Synapse software modulated the output of each LED and recorded photometry signals, which were sampled from the photodetector at 6.1 kHz. Demodulation of the brightness produced by the 465-nm excitation, which stimulates dopamine-dependent dLight fluorescence, versus isosbestic 415-nm excitation, which stimulates dLight in a dopamine-independent manner, allowed for correction for bleaching and movement artifacts.

Behavioral timestamps (e.g.; port entries) were fed into the RZ5P processor as TTL signals from the operant chambers (Med Associates) for alignment with neural data. Video recording feeds were similarly integrated with the photometry data via Synapse software. Photometry recordings occurred on the following sessions: habituation, 1, 3, 5, 6 (reversal 1), 8, 10, 11 (reversal 2), 13, 15, and 16 (extinction).

### DeepLabCut-based pose tracking

Markerless tracking of animal body parts was conducted using the DeepLabCut Toolbox (DLC version 2.2.3; (Mathis et al., 2018), and analysis of movement features based on these tracked coordinates was conducted in Matlab R2022b (Mathworks). All DLC analysis was conducted on either a Dell G7-7590 laptop running Windows 10 with an Intel Core i7-9750H CPU, 2.60Ghz, 16 GB RAM, and an NVIDIA GeForce RTX 2080 Max-Q 8 GB GPU or an Alienware Aurora R13 Gaming desktop running Windows 11 with an Intel Core i9-12900F CPU, 2400 Mhz, 128 GB RAM, and an NVIDIA GeForce RTX 3090 32 GB GPU. DeepLabCut was installed in an Anaconda environment with Python 3.8.15, CUDA 12.0 and Tensorflow 2.10.0. Videos (944 × 480 resolution) were recorded with a sampling frequency of 10-20 frames per second using TDT Synapse software with overhead cameras (Vanxse CCTV 960H 1000TVL HD Mini Spy Security Camera 2.8-12 mm Varifocal Lens Indoor Surveillance Camera).

### DeepLabCut model

2090 frames from 35 videos (32 different animals, 3 experiments) were labeled and 807 outlier frames were relabelled to refine a network described previously (Collins et al., 2023) for the current study. Labeled frames were split into a training set (95% of frames) and a test set (5% of frames). A ResNet-50 based neural network (Insafutdinov et al., 2016) was used for 1,030,000 training iterations. After final refinement we used a p-cutoff of 0.85 resulting in training set error of 2.99 pixels and test set error of 3.68 pixels. The body parts labeled included the nose, eyes, ears, fiber optic implant, shoulders, tail base, and an additional three points along the spine. Features of the environment were also labeled, including the 4 corners of the apparatus floor and the two magazine ports. This model was then used to analyze videos from 23 rats (10 DLS, 8 NAc core, and 5 NAc shell) for all training sessions.

### Histology

Rats received i.p. injections of Fatal-Plus (2 ml/kg; Patterson Veterinary) to induce a deep anesthesia, andwere transcardially perfused with cold phosphate buffered saline (PBS) followed by 4% paraformaldehyde (PFA). Brains were removed and post-fixed in 4% PFA for ~24 h, then cryoprotected in a 30% sucrose in PBS solution for at least 48 h. Sections were cut at 50 μm on a cryostat (Leica Biosystems). To confirm viral expression and optic fiber placements, brain sections containing the striatum were mounted on microscope slides and coverslipped with Vectashield containing DAPI counterstain. Slides were imaged using a fluorescent microscope (Keyence BZ-X710) with a 4x air immersion objective. Fiber placements and virus spreads were assessed using the Rat Brain Atlas (Paxinos and Watson, 2007).

### Behavioral analysis

Port entries and freezing bouts were the primary behavioral output measures. Port entry frequency and duration was analyzed from Med-PC software outputs and the generated TTL signals, while freezing bouts were calculated from positional information tracked using DLC. DLC coordinates and confidence values for each bodypart and frame were imported to Matlab and filtered to exclude body parts/features from any frame where the confidence was < 0.7. For labeled features of the environment (those with a fixed location), the average coordinates for that recording were used for analysis. For each video, a pixel-to-cm conversion rate was used to convert pixel distances to the real chamber dimensions. The distance (in pixels) between each edge of the environment floor and the diagonal measurements from corner to corner was measured, and these values were divided by the actual distance in cm. The mean of these values was then used as the conversion factor. The movement threshold for detecting freezing was calibrated to animal size using a scale factor determined from the relationship between body size (distance between the shoulder and lower back points) and the optimal threshold for detecting movement in a subset of animals used in this study (n = 4). This limit was used to detect movement in the face and head, and for the remaining body parts this value was multiplied by 2 to allow for detection of finer movements of the face vs the body. Freezing bouts were detected when all visible body parts were below their respective movement thresholds for a single frame, and a sliding window was used to determine when the speed of 2 or more body parts exceeded the detection threshold for 0.2 s, indicating the beginning and end of a movement. Frames in which < 2 body parts were visible were ignored and bouts with durations less than 1 s were excluded.

### Fiber Photometry Analysis

Recorded fluorescence signals produced by 465 nm and 415 nm LEDs were low pass filtered (2 Hz) and downsampled to 40 Hz, and a least-squares linear fit was applied to the 415 nm signal to align it to the 465 nm signal. The fitted 415 nm signal was used to normalize the 465 nm signal, where ΔF/F = (465 nm signal – fitted 415 nm signal)/(fitted 415 nm signal). Task events (e.g., cue presentations) were timestamped in the photometry data file via a TTL signal from Med-PC, and behavior was video recorded as described below at 10-20 FPS, with timestamps for each frame captured in the photometry file. Normalized signals for each trial were extracted in the window from 5 s preceding to 25 s following each cue presentation and Z-scored to the 5 s pre-cue period for each trial to minimize the effects of drift in the signal across experiment duration. Normalized signals for reward consumption were extracted in the window from 3 s preceding to 10 s following each port entry and Z-scored to the 3 s pre-port entry. A port entry was categorized as rewarded if it was the first port entry after reward delivery with a duration > 0.5 s. Normalized signals for shock responses were extracted in the window from 3 s preceding to 5 s following each shock delivery and Z-scored to the 3 s pre-shock delivery. Cue, reward, and shock responses were detected in the average waveform for each animal in the 2 s window beginning at event onset. The maximum and minimum responses in each window were detected, and latency was calculated relative to the start of the detection window. The slope of the rising and falling phase of the peak and trough waveforms were calculated between the maximum and 50% of max using the formula t2-t1/f2-f1, where t is the beginning and end timepoints of the rising or falling phase of the waveform, and f is the Z-scored fluorescence at each timepoint. Area under the curve (AUC) values were calculated from the Z-scored traces by numerical integration via the trapezoidal method using the trapz function (Matlab).

### Statistical analyses

Behavior and photometry signal data were analyzed with a combination of ANOVA models (one-way, twoway, and mixed effects). Post-hoc comparisons and planned t-tests were used to clarify main effects and interactions. Graphs represent average values of sites/ subjects, not individual trials. Data are expressed as mean ± s.e.m. For all tests, the α level for significance was set to p<0.05.

## SUPPLEMENTARY MATERIAL

**Supplementary Figure 1.**
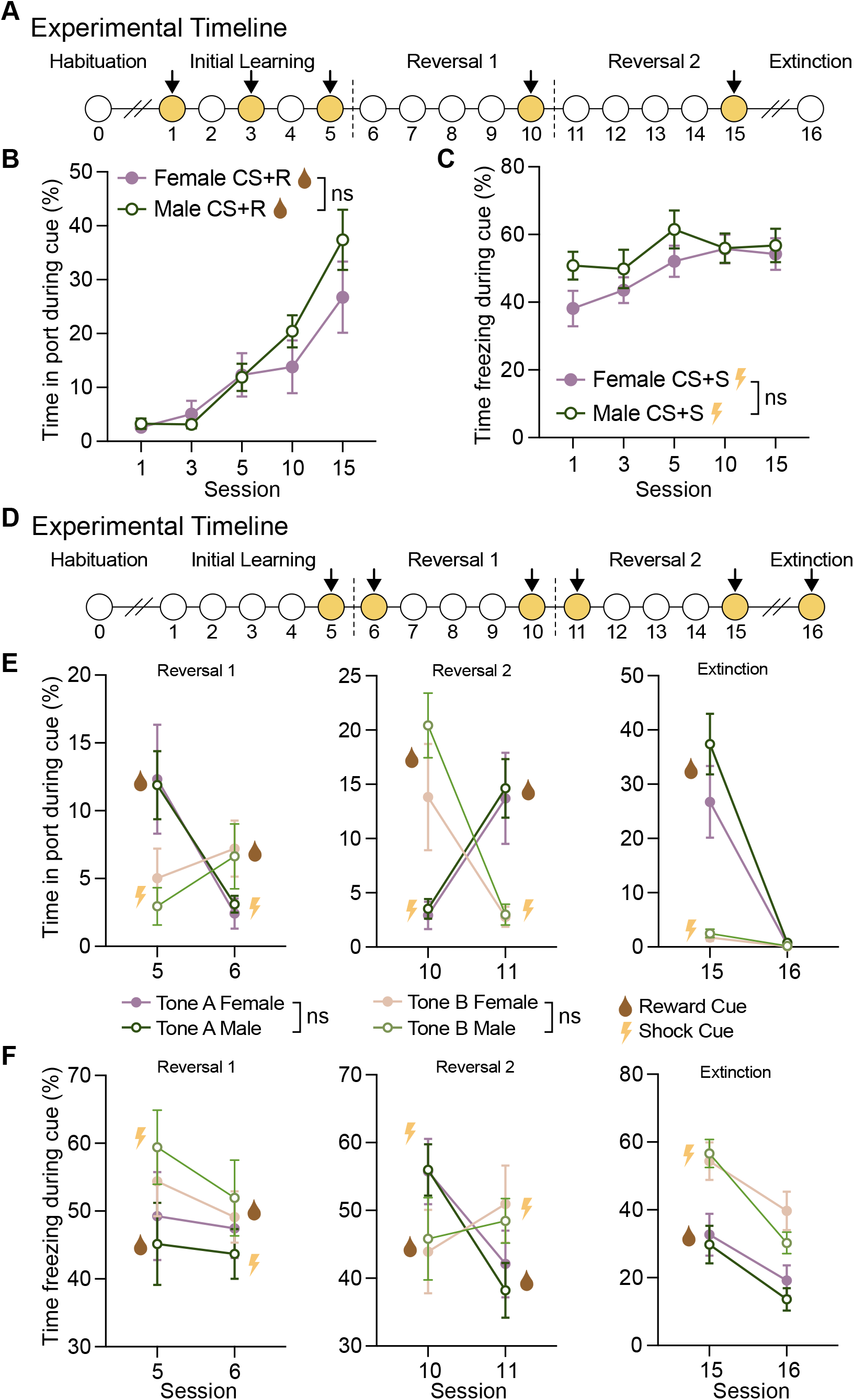
No sex differences in Pavlovian valence discrimination or reversal. A Experimental timeline showing the sessions highlighted in panels B and C. Female (n=11) and male (n=12) showed similar levels of B percent time spent in the reward port during reward cue presentations and C percent time spent freezing during shock cue presentations, across training. D Experimental timeline showing the sessions highlighted in panels E and F. Female and male rats showed similar levels of behavioral flexibility. E The percent time spent in the reward port during either cue was similar for males and females before and after reversal 1 (left), reversal 2 (middle), and extinction (right). F The percent time spent freezing during either cue was similar for males and females before and after reversal 1 (left), reversal 2 (middle), and extinction (right).

**Supplementary Figure 2.**
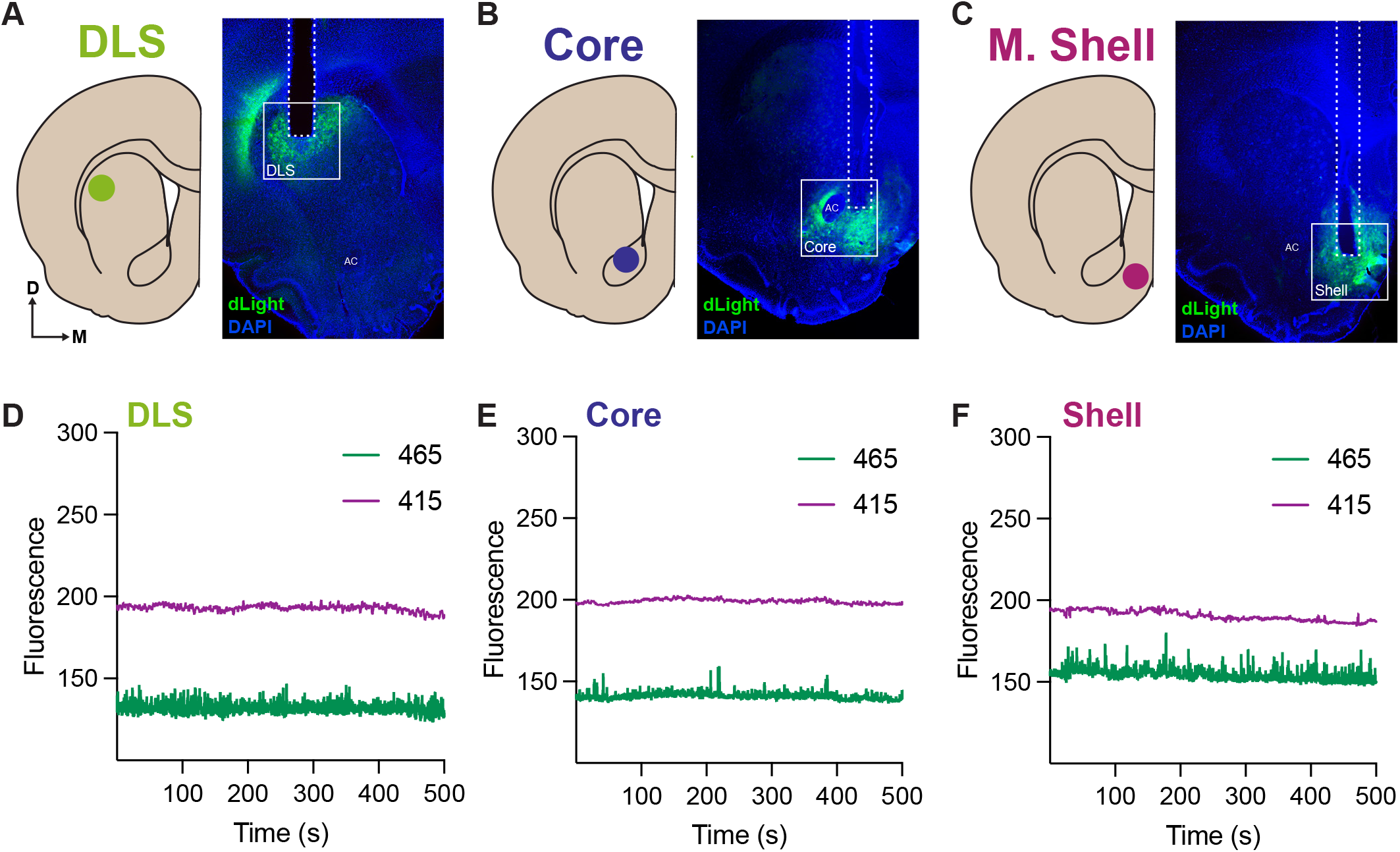
In vivo fiber photometry dopamine recordings across the striatum. Representative virus and fiber placement for the (A) DLS, (B) core, and (C) medial shell subregions. (D-F) Example longer (500 s) recording traces of raw 415 and 465 signals show consistent bleaching patterns across signal type and region.

**Supplementary Figure 3.**
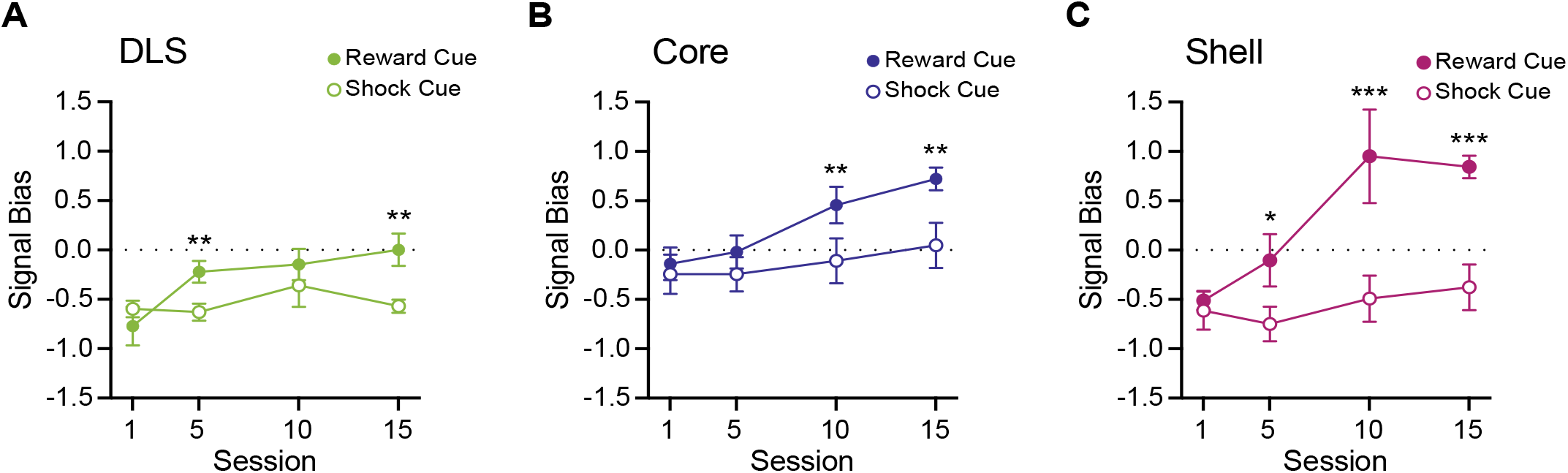
Dopamine signal response bias for reward versus shock cues. Dopamine response bias was computed by comparing the magnitude of the peak (most positive) and trough (most negative) parts of the signal following cue onset. (A-C) Signal bias for the reward and shock predictive CSs changed with different patterns across striatal subregions as training progressed. ***p<0.001, **p<0.01, *p<0.05.

**Supplementary Figure 4.**
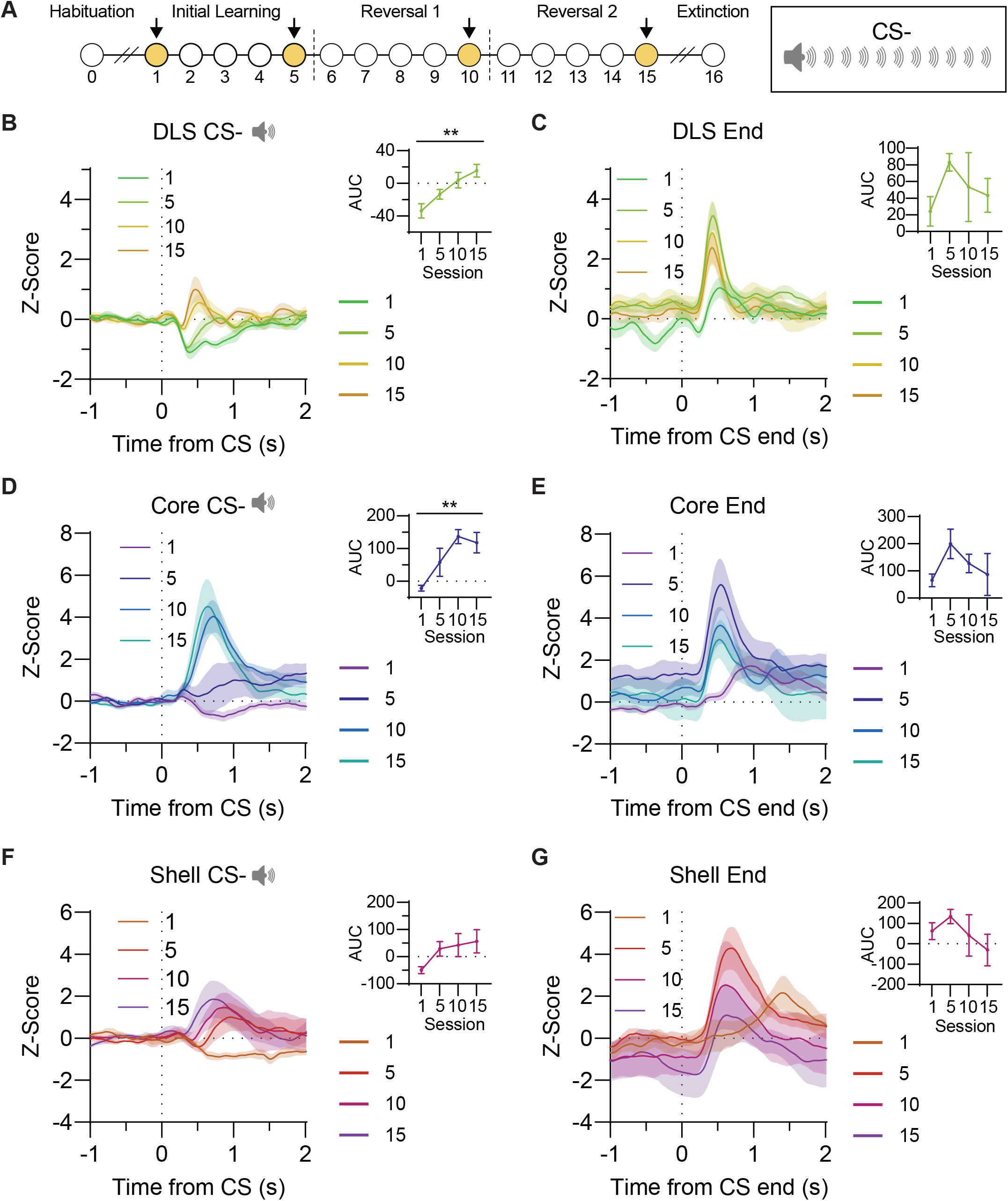
Robust dopamine signals emerge to the CS-in the multivalent cue discrimination task. A Experimental timeline. Photometry data from the highlighted sessions (1, 5, 10, 15) are shown in this figure for the CS-, presentations of which were intermingled with reward and shock-predictive cues. The CS-was never associated with either unconditioned stimulus. In the (B) DLS, (D) core, and (F) shell, dLight responses to the CS-increased across training. dLight responses were also seen following CS-offset in the (C) DLS, (E) core, and (G) shell, which generally grew in magnitude before stabilizing in the middle of training. All bars represent mean ± SEM. **p<0.01.

## REFERENCES

Badrinarayan, A., Wescott, S.A., Vander Weele, C.M., Saunders, B.T., Couturier, B.E., Maren, S., Aragona, B.J., 2012. Aversive stimuli differentially modulate real-time dopamine transmission dynamics within the nucleus accumbens core and shell. J. Neurosci. Off. J. Soc. Neurosci. 32, 15779–15790. 10.1523/JNEUROSCI.3557-12.2012

Bakhurin, K.I., Mac, V., Golshani, P., Masmanidis, S.C., 2016. Temporal correlations among functionally specialized striatal neural ensembles in reward-conditioned mice. J. Neurophysiol. 115, 1521–1532. 10.1152/jn.01037.2015

Barnes, T.D., Kubota, Y., Hu, D., Jin, D.Z., Graybiel, A.M., 2005. Activity of striatal neurons reflects dynamic encoding and recoding of procedural memories. Nature 437, 1158–1161. 10.1038/nature04053

Beste, C., Adelhöfer, N., Gohil, K., Passow, S., Roessner, V., Li, S.-C., 2018. Dopamine Modulates the Efficiency of Sensory Evidence Accumulation During Perceptual Decision Making. Int. J. Neuropsychopharmacol. 21, 649–655. 10.1093/ijnp/pyy019

Bromberg-Martin, E.S., Matsumoto, M., Hikosaka, O., 2010. Dopamine in motivational control: rewarding, aversive, and alerting. Neuron 68, 815–834. 10.1016/j.neu-ron.2010.11.022

Bromberg-Martin, E.S., Monosov, I.E., 2020. Neural circuitry of information seeking. Curr. Opin. Behav. Sci., Curiosity (Explore vs Exploit) 35, 62–70. 10.1016/j.cobe-ha.2020.07.006

Brown, H.D., McCutcheon, J.E., Cone, J.J., Ragozzino, M.E., Roitman, M.F., 2011. Primary food reward and reward-predictive stimuli evoke different patterns of phasic dopamine signaling throughout the striatum. Eur. J. Neurosci. 34, 1997–2006. 10.1111/j.1460-9568.2011.07914.x

Burgos-Robles, A., Kimchi, E.Y., Izadmehr, E.M., Porzenheim, M.J., Ramos-Guasp, W.A., Nieh, E.H., Felix-Ortiz, A.C., Namburi, P., Leppla, C.A., Presbrey, K.N., Anandalingam, K.K., Pagan-Rivera, P.A., Anahtar, M., Beyeler, A., Tye, K.M., 2017. Amygdala inputs to prefrontal cortex guide behavior amid conflicting cues of reward and punishment. Nat. Neurosci. 20, 824–835. 10.1038/nn.4553

Cardinal, R.N., Parkinson, J.A., Lachenal, G., Halkerston, K.M., Rudarakanchana, N., Hall, J., Morrison, C.H., Howes, S.R., Robbins, T.W., Everitt, B.J., 2002. Effects of selective excitotoxic lesions of the nucleus accumbens core, anterior cingulate cortex, and central nucleus of the amygdala on autoshaping performance in rats. Behav. Neurosci. 116, 553–567. 10.1037//0735-7044.116.4.553

Castañé, A., Theobald, D.E.H., Robbins, T.W., 2010. Selective lesions of the dorsomedial striatum impair serial spatial reversal learning in rats. Behav. Brain Res. 210, 74–83. 10.1016/j.bbr.2010.02.017

Collins, A.L., Saunders, B.T., 2020. Heterogeneity in striatal dopamine circuits: Form and function in dynamic reward seeking. J. Neurosci. Res. 98, 1046–1069. 10.1002/jnr.24587

Collins, V., Bornhoft, K.N., Wolff, A., Sinha, S., Saunders, B.T., 2023. Hierarchical cue control of cocaine seeking in the face of cost. Psychopharmacology (Berl.) 240, 461–476. 10.1007/s00213-022-06218-1

Cox, J., Witten, I.B., 2019. Striatal circuits for reward learning and decision-making. Nat. Rev. Neurosci. 20, 482–494. 10.1038/s41583-019-0189-2

Day, H.L.L., Reed, M.M., Stevenson, C.W., 2016. Sex differences in discriminating between cues predicting threat and safety. Neurobiol. Learn. Mem. 133, 196–203. 10.1016/j.nlm.2016.07.014

De Jong, J.W., Afjei, S.A., Pollak Dorocic, I., Peck, J.R., Liu, C., Kim, C.K., Tian, L., Deisseroth, K., Lammel, S., 2019. A Neural Circuit Mechanism for Encoding Aversive Stimuli in the Mesolimbic Dopamine System. Neuron 101, 133-151.e7. 10.1016/j.neuron.2018.11.005

Di Ciano, P., Cardinal, R.N., Cowell, R.A., Little, S.J., Everitt, B.J., 2001. Differential involvement of NMDA, AMPA/kainate, and dopamine receptors in the nucleus accumbens core in the acquisition and performance of pavlovian approach behavior. J. Neurosci. Off. J. Soc. Neurosci. 21, 9471–9477. 10.1523/JNEUROSCI.21-23-09471.2001

Everitt, B.J., Robbins, T.W., 2005. Neural systems of reinforcement for drug addiction: from actions to habits to compulsion. Nat. Neurosci. 8, 1481–1489. 10.1038/nn1579

Fadok, J.P., Dickerson, T.M.K., Palmiter, R.D., 2009. Dopamine Is Necessary for Cue-Dependent Fear Conditioning. J. Neurosci. 29, 11089–11097. 10.1523/JNEUROS-CI.1616-09.2009

Fiorillo, C.D., Tobler, P.N., Schultz, W., 2003. Discrete Coding of Reward Probability and Uncertainty by Dopamine Neurons. Science 299, 1898–1902. 10.1126/science.1077349

Fraser, K.M., Collins, V.L., Wolff, A.R., Ottenheimer, D.J., Bornhoft, K.N., Pat, F., Chen, B.J., Janak, P.H., Saunders, B.T., 2023. Contexts facilitate dynamic value encoding in the mesolimbic dopamine system. BioRxiv Prepr. Serv. Biol. 2023.11.05.565687. 10.1101/2023.11.05.565687

Goedhoop, J.N., van den Boom, B.J., Robke, R., Veen, F., Fellinger, L., van Elzelingen, W., Arbab, T., Willuhn, I., 2022. Nucleus accumbens dopamine tracks aversive stimulus duration and prediction but not value or prediction error. eLife 11, e82711. 10.7554/eLife.82711

Gottlieb, J., Hayhoe, M., Hikosaka, O., Rangel, A., 2014. Attention, Reward, and Information Seeking. J. Neurosci. 34, 15497–15504. 10.1523/JNEUROSCI.3270-14.2014

Haluk, D.M., Floresco, S.B., 2009. Ventral Striatal Dopamine Modulation of Different Forms of Behavioral Flexibility. Neuropsychopharmacology 34, 2041–2052. 10.1038/npp.2009.21

Hart, A.S., Clark, J.J., Phillips, P.E.M., 2015. Dynamic shaping of dopamine signals during probabilistic Pavlovian conditioning. Neurobiol. Learn. Mem., Memory and decision making 117, 84–92. 10.1016/j.nlm.2014.07.010

Horvitz, J.C., 2000. Mesolimbocortical and nigrostriatal dopamine responses to salient non-reward events. Neuroscience 96, 651–656. 10.1016/S0306-4522(00)00019-1

Howe, M.W., Dombeck, D.A., 2016. Rapid signalling in distinct dopaminergic axons during locomotion and reward. Nature 535, 505–510. 10.1038/nature18942

Izquierdo, A., Jentsch, J.D., 2012. Reversal learning as a measure of impulsive and compulsive behavior in addictions. Psychopharmacology (Berl.) 219, 607–620. 10.1007/s00213-011-2579-7

Kalmbach, A., Winiger, V., Jeong, N., Asok, A., Gallistel, C.R., Balsam, P.D., Simpson, E.H., 2022. Dopamine encodes real-time reward availability and transitions between reward availability states on different timescales. Nat. Commun. 13, 3805. 10.1038/s41467-022-31377-2

Keiflin, R., Pribut, H.J., Shah, N.B., Janak, P.H., 2019. Ventral Tegmental Dopamine Neurons Participate in Reward Identity Predictions. Curr. Biol. 29, 93-103.e3. 10.1016/j.cub.2018.11.050

Kim, H.F., Ghazizadeh, A., Hikosaka, O., 2015. Dopamine Neurons Encoding Long-Term Memory of Object Value for Habitual Behavior. Cell 163, 1165–1175. 10.1016/j.cell.2015.10.063

Klanker, M., Feenstra, M., Denys, D., 2013. Dopaminergic control of cognitive flexibility in humans and animals. Front. Neurosci. 7. 10.3389/fnins.2013.00201

Klanker, M., Fellinger, L., Feenstra, M., Willuhn, I., Denys, D., 2017. Regionally distinct phasic dopamine release patterns in the striatum during reversal learning. Neuroscience, Cognitive Flexibility: Development, Disease, and Treatment 345, 110–123. 10.1016/j.neuroscience.2016.05.011

Kutlu, M.G., Tat, J., Christensen, B.A., Zachry, J.E., Calipari, E.S., 2023. Dopamine release at the time of a predicted aversive outcome causally controls the trajectory and expression of conditioned behavior. Cell Rep. 42, 112948. 10.1016/j.celrep.2023.112948

Kutlu, M.G., Zachry, J.E., Brady, L.J., Melugin, P.R., Kelly, S.J., Sanders, C., Tat, J., Johnson, A.R., Thibeault, K., Lopez, A.J., Siciliano, C.A., Calipari, E.S., 2020. A novel multidimensional reinforcement task in mice elucidates sex-specific behavioral strategies. Neuropsychopharmacology 45, 1463–1472. 10.1038/s41386-020-0692-1

Kutlu, M.G., Zachry, J.E., Melugin, P.R., Cajigas, S.A., Chevee, M.F., Kelly, S.J., Kutlu, B., Tian, L., Siciliano, C.A., Calipari, E.S., 2021. Dopamine release in the nucleus accumbens core signals perceived saliency. Curr. Biol. 31, 4748-4761.e8. 10.1016/j.cub.2021.08.052

Lak, A., Nomoto, K., Keramati, M., Sakagami, M., Kepecs, A., 2017. Midbrain Dopamine Neurons Signal Belief in Choice Accuracy during a Perceptual Decision. Curr. Biol. 27, 821–832. 10.1016/j.cub.2017.02.026

Lak, A., Okun, M., Moss, M.M., Gurnani, H., Farrell, K., Wells, M.J., Reddy, C.B., Kepecs, A., Harris, K.D., Carandini, M., 2020. Dopaminergic and Prefrontal Basis of Learning from Sensory Confidence and Reward Value. Neuron 105, 700-711.e6. 10.1016/j.neuron.2019.11.018

Lammel, S., Lim, B.K., Malenka, R.C., 2014. Reward and aversion in a heterogeneous midbrain dopamine system. Neuropharmacology, NIDA 40th Anniversary Issue 76, 351–359. 10.1016/j.neuropharm.2013.03.019

Leeson, V.C., Robbins, T.W., Matheson, E., Hutton, S.B., Ron, M.A., Barnes, T.R.E., Joyce, E.M., 2009. Discrimination Learning, Reversal, and Set-Shifting in First-Episode Schizophrenia: Stability Over Six Years and Specific Associations with Medication Type and Disorganization Syndrome. Biol. Psychiatry, Cortical Development and Glutamatergic Dysregulation in Schizophrenia 66, 586–593. 10.1016/j.biopsych.2009.05.016

Lerner, T.N., 2020. Interfacing behavioral and neural circuit models for habit formation. J. Neurosci. Res. 98, 1031–1045. 10.1002/jnr.24581

Lerner, T.N., Shilyansky, C., Davidson, T.J., Evans, K.E., Beier, K.T., Zalocusky, K.A., Crow, A.K., Malenka, R.C., Luo, L., Tomer, R., Deisseroth, K., 2015. Intact-Brain Analyses Reveal Distinct Information Carried by SNc Dopamine Subcircuits. Cell 162, 635–647. 10.1016/j.cell.2015.07.014

Luo, R., Uematsu, A., Weitemier, A., Aquili, L., Koivumaa, J., McHugh, T.J., Johansen, J.P., 2018. A dopaminergic switch for fear to safety transitions. Nat. Commun. 9, 2483. 10.1038/s41467-018-04784-7

Malvaez, M., Wassum, K.M., 2018. Regulation of habit formation in the dorsal striatum. Curr. Opin. Behav. Sci., Habits and Skills 20, 67–74. 10.1016/j.cobeha.2017.11.005

Mathis, A., Mamidanna, P., Cury, K.M., Abe, T., Murthy, V.N., Mathis, M.W., Bethge, M., 2018. DeepLabCut: markerless pose estimation of user-defined body parts with deep learning. Nat. Neurosci. 21, 1281–1289. 10.1038/s41593-018-0209-y

Matsumoto, M., Hikosaka, O., 2009. Two types of dopamine neuron distinctly convey positive and negative motivational signals. Nature 459, 837–841. 10.1038/nature08028

Mccutcheon, J.E., Ebner, S.R., Loriaux, A.L., Roitman, M.F., 2012. Encoding of Aversion by Dopamine and the Nucleus Accumbens. Front. Neurosci. 6. 10.3389/fnins.2012.00137

Mikhael, J.G., Kim, H.R., Uchida, N., Gershman, S.J., 2022. The role of state uncertainty in the dynamics of dopamine. Curr. Biol. 32, 1077-1087.e9. 10.1016/j.cub.2022.01.025

Mohebi, A., Wei, W., Pelattini, L., Kim, K., Berke, J.D., 2024. Dopamine transients follow a striatal gradient of reward time horizons. Nat. Neurosci. 27, 737–746. 10.1038/s41593-023-01566-3

Nestler, E.J., Lüscher, C., 2019. The Molecular Basis of Drug Addiction: Linking Epigenetic to Synaptic and Circuit Mechanisms. Neuron 102, 48–59. 10.1016/j.neu-ron.2019.01.016

Ng, K.H., Pollock, M.W., Urbanczyk, P.J., Sangha, S., 2018. Altering D1 receptor activity in the basolateral amygdala impairs fear suppression during a safety cue. Neurobiol. Learn. Mem. 147, 26–34. 10.1016/j.nlm.2017.11.011

Ng, K.H., Sangha, S., 2023. Encoding of conditioned inhibitors of fear in the infralimbic cortex. Cereb. Cortex 33, 5658–5670. 10.1093/cercor/bhac450

Oleson, E.B., Gentry, R.N., Chioma, V.C., Cheer, J.F., 2012. Subsecond dopamine release in the nucleus accumbens predicts conditioned punishment and its successful avoidance. J. Neurosci. Off. J. Soc. Neurosci. 32, 14804–14808. 10.1523/JNEUROSCI.3087-12.2012

Parker, N.F., Cameron, C.M., Taliaferro, J.P., Lee, J., Choi, J.Y., Davidson, T.J., Daw, N.D., Witten, I.B., 2016. Reward and choice encoding in terminals of midbrain dopamine neurons depends on striatal target. Nat. Neurosci. 19, 845–854. 10.1038/nn.4287

Paxinos, G., Watson, C., 2007. The Rat Brain in Stereotaxic Coordinates, 6th ed. Elsevier Inc.

Pezze, M.A., Feldon, J., 2004. Mesolimbic dopaminergic pathways in fear conditioning. Prog. Neurobiol. 74, 301–320. 10.1016/j.pneurobio.2004.09.004

Radke, A.K., Kocharian, A., Covey, D.P., Lovinger, D.M., Cheer, J.F., Mateo, Y., Holmes, A., 2019. Contributions of nucleus accumbens dopamine to cognitive flexibility. Eur. J. Neurosci. 50, 2023–2035. 10.1111/ejn.14152

Ragozzino, M.E., 2007. The Contribution of the Medial Prefrontal Cortex, Orbitofrontal Cortex, and Dorsomedial Striatum to Behavioral Flexibility. Ann. N. Y. Acad. Sci. 1121, 355–375. 10.1196/annals.1401.013

Redgrave, P., Prescott, T.J., Gurney, K., 1999. Is the short-latency dopamine response too short to signal reward error? Trends Neurosci. 22, 146–151. 10.1016/S0166-2236(98)01373-3

Remijnse, P.L., Nielen, M.M.A., van Balkom, A.J.L.M., Cath, D.C., van Oppen, P., Uylings, H.B.M., Veltman, D.J., 2006. Reduced orbitofrontal-striatal activity on a reversal learning task in obsessive-compulsive disorder. Arch. Gen. Psychiatry 63, 1225–1236. 10.1001/archpsyc.63.11.1225

Saunders, B.T., Richard, J.M., Margolis, E.B., Janak, P.H., 2018. Dopamine neurons create Pavlovian conditioned stimuli with circuit-defined motivational properties. Nat. Neurosci. 21, 1072–1083. 10.1038/s41593-018-0191-4

Saunders, B.T., Robinson, T.E., 2012. The role of dopamine in the accumbens core in the expression of Pavlovian-conditioned responses. Eur. J. Neurosci. 36, 2521–2532. 10.1111/j.1460-9568.2012.08217.x

Schultz, W., Dayan, P., Montague, P.R., 1997. A Neural Substrate of Prediction and Reward. Science 275, 1593–1599. 10.1126/science.275.5306.1593

Sharpe, M.J., Chang, C.Y., Liu, M.A., Batchelor, H.M., Mueller, L.E., Jones, J.L., Niv, Y., Schoenbaum, G., 2017. Dopamine transients are sufficient and necessary for acquisition of model-based associations. Nat. Neurosci. 20, 735–742. 10.1038/nn.4538

Stelly, C.E., Haug, G.C., Fonzi, K.M., Garcia, M.A., Tritley, S.C., Magnon, A.P., Ramos, M.A.P., Wanat, M.J., 2019. Pattern of dopamine signaling during aversive events predicts active avoidance learning. Proc. Natl. Acad. Sci. U. S. A. 116, 13641–13650. 10.1073/pnas.1904249116

Swainson, R., Rogers, R.D., Sahakian, B.J., Summers, B.A., Polkey, C.E., Robbins, T.W., 2000. Probabilistic learning and reversal deficits in patients with Parkinson’s disease or frontal or temporal lobe lesions: possible adverse effects of dopaminergic medication. Neuropsychologia 38, 596–612. 10.1016/s0028-3932(99)00103-7

Takahashi, Y.K., Batchelor, H.M., Liu, B., Khanna, A., Morales, M., Schoenbaum, G., 2017. Dopamine Neurons Respond to Errors in the Prediction of Sensory Features of Expected Rewards. Neuron 95, 1395-1405.e3. 10.1016/j.neu-ron.2017.08.025

Ungless, M.A., 2004. Dopamine: the salient issue. Trends Neurosci. 27, 702–706. 10.1016/j.tins.2004.10.001

van Elzelingen, W., Goedhoop, J., Warnaar, P., Denys, D., Arbab, T., Willuhn, I., 2022. A unidirectional but not uniform striatal landscape of dopamine signaling for motivational stimuli. Proc. Natl. Acad. Sci. U. S. A. 119, e2117270119. 10.1073/pnas.2117270119

Vellani, V., de Vries, L.P., Gaule, A., Sharot, T., 2020. A selective effect of dopamine on information-seeking. eLife 9, e59152. 10.7554/eLife.59152

Ventura, R., Morrone, C., Puglisi-Allegra, S., 2007. Prefrontal/accumbal catecholamine system determines motivational salience attribution to both reward- and aversion-related stimuli. Proc. Natl. Acad. Sci. U. S. A. 104, 5181–5186. 10.1073/pnas.0610178104

Wendler, E., Gaspar, J.C.C., Ferreira, T.L., Barbiero, J.K., Andreatini, R., Vital, M.A.B.F., Blaha, C.D., Winn, P., Da Cunha, C., 2014. The roles of the nucleus accumbens core, dorsomedial striatum, and dorsolateral striatum in learning: Performance and extinction of Pavlovian fear-conditioned responses and instrumental avoidance responses. Neurobiol. Learn. Mem. 109, 27–36. 10.1016/j.nlm.2013.11.009

Wise, R.A., 2004. Dopamine, learning and motivation. Nat. Rev. Neurosci. 5, 483–494. 10.1038/nrn1406

Yin, H.H., Knowlton, B.J., 2006. The role of the basal ganglia in habit formation. Nat. Rev. Neurosci. 7, 464–476. 10.1038/nrn1919

Yin, H.H., Mulcare, S.P., Hilário, M.R.F., Clouse, E., Holloway, T., Davis, M.I., Hansson, A.C., Lovinger, D.M., Costa, R.M., 2009. Dynamic reorganization of striatal circuits during the acquisition and consolidation of a skill. Nat. Neurosci. 12, 333–341. 10.1038/nn.2261

